# Chronic opioid treatment arrests neurodevelopment and alters synaptic activity in human midbrain organoids

**DOI:** 10.1101/2021.06.02.446827

**Authors:** Hye Sung Kim, Yang Xiao, Xuejing Chen, Siyu He, Jongwon Im, Moshe J. Willner, Michael O. Finlayson, Cong Xu, Huixiang Zhu, Se Joon Choi, Eugene V. Mosharov, Hae-Won Kim, Bin Xu, Kam W. Leong

## Abstract

The impact of long-term opioid exposure on the embryonic brain is crucial to healthcare due to the surging number of pregnant mothers with an opioid dependency. Current studies on the neuronal effects are limited due to human brain inaccessibility and cross-species differences among animal models. Here, we report a model to assess cell-type specific responses to acute and chronic fentanyl treatment, as well as fentanyl withdrawal, using human induced pluripotent stem cell (hiPSC)-derived midbrain organoids. Single cell mRNA sequencing (25,510 single cells in total) results suggest that chronic fentanyl treatment arrests neuronal subtype specification during early midbrain development and alters the pathways associated with synaptic activities and neuron projection. Acute fentanyl treatment, however, increases dopamine release but does not induce significant changes in gene expressions of cell lineage development. To date, our study is the first unbiased examination of midbrain transcriptomics with synthetic opioid treatment at the single cell level.

## Introduction

The opioid epidemic has reached crisis levels across the globe, with opioid use disorder (OUD) affecting 40.5 million people worldwide^1^. The perverse misuse of prescription opioids and heroin combined with the emergence of extremely potent fentanyl derivatives triggered a 10-fold increase in overdose fatalities in the United states from 2013 to 2018^2–5^. Indeed, the negative social, economic, and health ramifications of COVID-19 have further worsened this situation, with significant increases in opioid-related use, misuse, and non-fatal/fatal overdose after the start of the pandemic^6, 7^. An unfortunate consequence of this accelerated opioid use includes a parallel surge in the use of prescription opioids among pregnant women^8, 9^. Survey data found that 6.6% of women reported prescription opioid use during pregnancy, with 21.2% of those women reporting misuse, and 22–30% of women filling at least one prescription for an opioid analgesic during pregnancy^9, 10^. In addition to increased health care costs and adverse maternal outcomes, opioid use during pregnancy leads to an inevitable increase in the incidence of neonates exposed to opioids *in utero*^11–13^.

Prenatal opioid exposure, which includes both the use and misuse of prescription and illicit opioid drugs, causes unusual and deleterious symptoms in neonates including neonatal abstinence syndrome (NAS), small head circumference^14^, decreased cerebral volume^15^, microstructural brain injury^9^, and low birth weight^8, 9, 16^. Extensive clinical data also suggest that *in utero* opioid exposure has adverse effects on the developing organ systems, including the central nervous system^9, 17, 18^. The impaired neurodevelopment of these neonates are correlated with long-term issues, with follow-up studies revealing that individuals who had *in utero* opioid exposure experience problems with cognitive, behavioral, and developmental outcomes in their childhood^19, 20^. However, the precise cellular and molecular mechanisms underlying opioid-related neurodevelopmental disruptions in humans are unclear, and a better understanding of potential risks of long-term opioid exposure is crucial. This is especially urgent because synthetic opioids (i.e. fentanyl, methadone, and buprenorphine) themselves are frequently used to treat both pregnant patients with OUD and neonates with NAS, and thus advancements in therapeutics are absolutely necessary^21, 22^.

Efforts to understand the mechanisms underlying human neurodevelopment in response to opioids are fraught with challenges, such as limited accessibility to fetal brain specimens, ethical conflicts, and complex individual drug abuse history. Although pre-clinical studies conducted with animal models have revealed the detrimental effects of opioid use on the neurodevelopment of offspring^23^, inherent species differences have led to a general gap in neuropsychiatric clinical translation and a subsequent failure in drug development^24^. Pharmacological therapeutics developed using animal models have also been particularly ineffective for the treatment of opioid abuse in humans because: 1) different species vary in their neurodevelopmental trajectories, receptor expression, and central opioid pharmacokinetics, and 2) animal models cannot capture the broader network of symptoms and environmental/social factors that are fundamental to drug abuse and addiction^25, 26^. Moreover, although human postmortem brain specimens have the ability to directly assess the human-specific differences in gene expression, biomarkers, and neuroanatomy in drug abuse^27–30^, cellular and molecular responses cannot be evaluated in a time-dependent manner.

Brain organoids are thus excellent potential candidates to bridge the gap between animal models and human studies. Brain organoids are self-assembled three-dimensional aggregates generated from human pluripotent stem cells *in vitro* that resemble the human fetal brain^31^. Unlike conventional two-dimensional cell cultures, organoid models better recapitulate the fetal brain in cell type composition, 3D architecture, and similar lineage trajectory^32–35^. Brain organoids also provide a platform for unparalleled manipulation, enabling systematic studies of human neurodevelopment, disease modeling, and drug screening^33, 36–40^. Further, recent advances in guided organoid generation methods using lineage-specific patterning factors allow for the generation of micro-tissue mimicking certain brain regions such as the cerebral cortex^41, 42^, hippocampus^43^, and midbrain^36^.

In addition, brain organoids have revolutionized the characterization of brain development in combination with patient-derived iPSC engineering, genetic engineering, epigenetics, and single cell multiomics^44–47^. Specifically, single-cell RNA sequencing (scRNA-seq) offers insights into underlying molecular signatures. Systematic RNA-seq analysis could identify early events of human neurogenesis and synaptogenesis that are limited to experimental access and validate cell types and neuronal activities^38, 48, 49^. For example, recent research identified 545 differentially expressed genes in the midbrains of opioid users compared to non-opioid users using bulk RNA-seq^27^. To date, however, cellular changes observed in distinct human neuronal subtypes upon opioid exposure have not been reported. Although multiple tested brain organoid generation models have been developed and validated, functional models are yet to be established to explore the temporal and spatial changes of neurons in response to a specific stressor, such as opioids. The susceptibility of newborns and adults to develop altered opioid responses thus remains under-investigated and poorly understood.

To our knowledge, this is the first study to investigate the influences of chronic opioid exposure on molecular and cellular alterations during neurodevelopment using a human midbrain organoid model. Based on a previously reported organoid generation protocol^36^, we established our midbrain organoids as a viable model to study neurodevelopment, as well as developed a framework for evaluating cell type-specific transcriptomic responses to fentanyl exposure in these organoids. Moreover, our comprehensive molecular characterization shows pseudo-time metrics of midbrain organoid development and offers an in-depth analysis of affected gene features and pathways under a chronic fentanyl exposure condition.

Interestingly, chronic fentanyl treatment arrests ectodermal cells at the stage of neural progenitors or neuroblasts from progressing into neurons, while acute fentanyl treatment does not cause significant transcriptomic changes associated with neurodevelopment. In addition, although less neurons were observed in the fentanyl-treated organoids, the average gene expression level of each neuron is higher in neuron projection, synapse assembly, and neurotransmitter dynamics. Furthermore, neurodevelopment resumed in lineage-related gene expression when fentanyl was withdrawn. Our investigation into the arrested neurodevelopment in fentanyl-treated organoids, in addition to further advancements in organoid technology, can help inform improved therapeutics for opioid abuse and prenatal opioid exposure.

## Results

### Generation and characterization of midbrain organoids

We generated midbrain organoids using a protocol adapted from Kriks *et al*. and Jo *et al*. with minor modifications (**Figure 1A**, **Methods**)^36, 37^. Human iPSCs were dissociated to single cells to form uniformly-sized embryoid bodies (EBs) approximately 300μm in diameter in non-treated U-bottom 96-well plates (**Figure 1B**). At day 7-14, EBs showed uniform neuroectoderm formation along the outer surface of EBs where optically translucent and radially organized (**Figure 1B** (inset)). These neuroectoderm-containing spheroids were cultured in N2 neuronal media supplemented with neurotrophic factors (BDNF, Ascorbic acid, GDNF, cAMP, TGF-b3, and DAPT) once neural induction was achieved in the first 9 days (**Figure 1A**, **Methods**). The organoids grew up to 0.8~1.2mm in diameter after 21 days (**Figure 1C**). The cytoarchitecture of the organoids were examined at day 21 by immunocytochemical analysis (**Figure 1D**). The midbrain progenitors expressing OTX2 or FOXA2 proteins were located closer to the core of the organoids, with OTX2 expression specifically found in the apical surface and extending to the intermediate region of the neuroepithelia (**Figure 1D** and **Figure S1B**). In contrast, dopaminergic (TH positive) and GABAergic (GABA positive) neurons were detected along the outer edge of the organoids, which is similar to the layering of the human midbrain development (**Figure S1A**)^50^.

**Figure 1.**
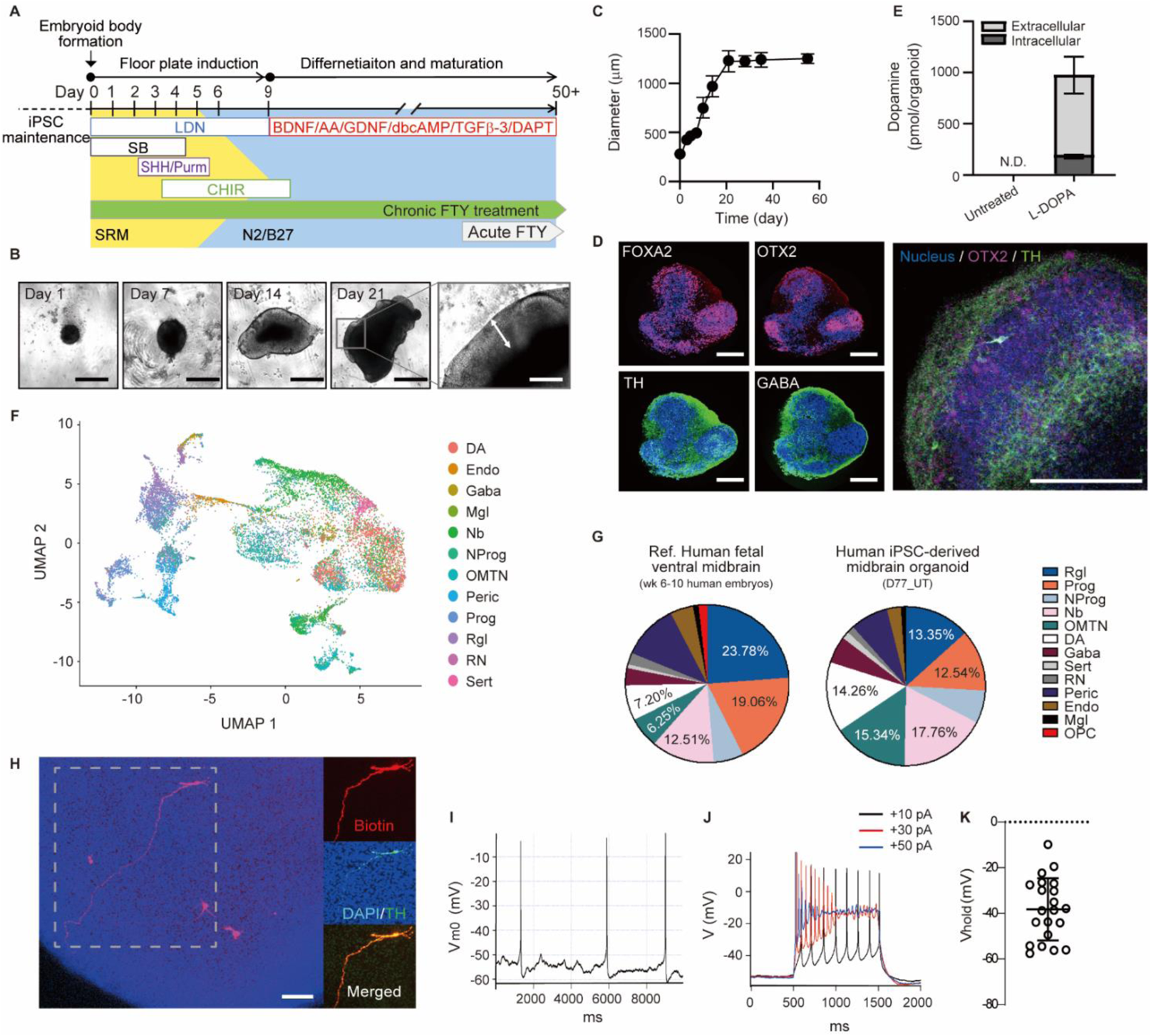
Generation and characterization of midbrain-like organoids. (A) Timeline of the midbrain-like organoid development. (B) Morphological changes and (C) growth (n=20 organoids, three individual experiments) of organoids over time (scale bars: 500 μm for day 1 - day 21, 200 μm for the inset). (D) Cytoarchitecture of the organoids examined at day 21 (scale bar, 200 μm). (E) Dopamine content in Day 35 organoids before and after 1h treatment with 100 µM L-DOPA (n=3 organoids). (F) Cell type identification and (G) compositions of all samples (Day 53 and Day 77 organoids). (H-K) Electrophysiological properties of neurons in Day 35 organoids. (H) Recorded cells were labeled with biotin dye (red) added to the patch pipette. After fixation, organoids were immunostained for TH (green). All recorded neurons (n=21) were confirmed to be TH+ (scale bar, 100μm). (I) Representative whole-cell recording from spontaneously active DA neuron. (J) Evoked action potentials following step current injections of +10pA (black), +30pA (red), and +50pA (blue). (K) Average resting membrane potential of DA neurons in organoids.

To further confirm whether organoids specifically developed towards the midbrain specification, we monitored the time course of a set of marker genes for 35 days by RT-PCR (**Figure S1C**). First, the expression of pluripotency markers, OCT4 and NANOG, drastically decreased upon neuronal induction, while pan-neuronal marker expression, TUJ1 and MAP2, gradually increased. In contrast, the expression of OTX2, a homeodomain transcription factor required for patterning the midbrain region^51^, increased after 7 days of floor plate induction. The co-expression of the floor-plate (FP) marker FOXA2 and the roof plate marker LMX1A, a unique feature of midbrain development^37^, was also observed. Furthermore, we observed robust expression of various midbrain specific neuronal markers. For example, the expression of midbrain dopaminergic neuronal markers (TH, DAT and PITX3) was up-regulated during the differentiation process. On the contrary, expression levels of PAX6, a forebrain marker, and TBR2 and GBX2, hindbrain markers, were low and not significantly changed. The immunocytochemical staining results also exhibited a decrease in the number of OTX2+ cells and an increase in the number of tyrosine hydroxylase (TH+) from day 7 to day 35 of the differentiation (**Figure S1A**). These results indicate that cells of the midbrain organoids gradually transitioned from proliferating neuroprogenitors into more mature midbrain-specific neurons.

To comprehensively determine the cell composition of the iPSC-derived midbrain organoids, single cell RNA sequencing was performed and annotated according to the cell type signatures of human fetal ventral midbrain (6-11 weeks human embryos) previously reported by Manno *et al*.^52^ Cells at two differential time points, day 53 and day 77, were analyzed. At each time point, cells from three organoids of each condition were pooled. A total of 25,510 single cells was analyzed (**Table S1**). We identified twelve cell types in the midbrain organoids including progenitor types (progenitors (Prog), radial glia-like cells (Rgl), and neural progenitors (NProg)), neuroblasts (Nb), neuronal cell types (oculomotor and trochlear nucleus (OMTN), Dopaminergic (DA), GABAergic (Gaba), red nucleus (RN), and Serotoninergic (Sert) neurons), and mesoderm-derived cell types (endothelial cells (Endo), pericytes (Peric), microglia-like cells (Mgl)) (**Figure 1F** and **G**, and **Figure S2, Methods**). Compared to human fetal midbrain, our organoids did not have oligodendrocyte progenitor cells (OPC) as of day 77 in culture. Except for OPC, overall cell type composition of the organoids was comparable to that of the human fetal ventral midbrain, with some variation in the proportions of each cell type (day 53 or 77 of *in vitro* differentiation vs. week 6-11 of *in vivo* embryonic development). From day 53 to day 77, the percentage of neuroblast and progenitor types decreased from 59% to 50%, whereas the percentage of neuronal and mesoderm-derived cell types increased from 35% to 38% and 6% to 12%, respectively (**Figure S2A**). This result further indicates the subsequent differentiation of neuroprogenitors into neurons in the developing organoids.

To determine the electrophysiological properties of the cells, at day 35 we immobilized the organoids on a Matrigel-coated coverslip and performed whole-cell current-clamp recordings from cells on or near the surface of the organoid. Biocytin dye was added to the patch pipette allowing to confirm in post-fixed organoids that recorded cells were TH+ (**Figure 1H**). Electrophysiological analysis revealed that ~20% of the recorded neurons (4 of 21 cells) fired spontaneous action potentials with an average frequency of 0.6 ± 0.3 Hz (**Figure 1I**), which is lower than the ~4 Hz autonomous firing frequency of mature mouse DA neurons. Injection of a +10 pA step current (**Figure 1J**, black) produced a train of stable action potentials, but higher current steps (+30 pA (red) and +50 pA (blue)) yielded action potentials with a higher frequency and lower amplitude that quickly ceased. These results suggest that the dopaminergic neurons of the organoid are not fully matured at day 35 of differentiation. Further, the average resting membrane potential of the neurons in the organoids was around −40mV, which is higher than that of mature neurons (~−60 mV) (**Figure 1K**). Similarly, dopamine synthesis and release were undetectable in untreated organoids, but increased rapidly in the presence of dopamine precursor L-dihydroxyphenylalanine (L-DOPA), demonstrating the ability of neurons to produce the neurotransmitter (**Figure 1E**). Taken together, these results suggest that midbrain organoids are similar to early-stage human fetal midbrains and could thus be used as a tool for understanding the influence of chronic opioid exposure on neurodevelopment.

### Cellular responses to acute opioid exposure for 4 hours

First, we confirmed that organoids express opioid receptors, including mu (OPRM1), kappa (OPRK1), and delta (OPRD1). The mRNA expression levels of opioid receptors drastically increased until day 35 and saturated after that point (**Figure 2A**). In addition, the opioid receptors were detected where TH+ or GABA+ neurons were located (**Figure 2B**), indicating that the midbrain-specific neurons of the organoid express opioid receptors.

**Figure 2.**
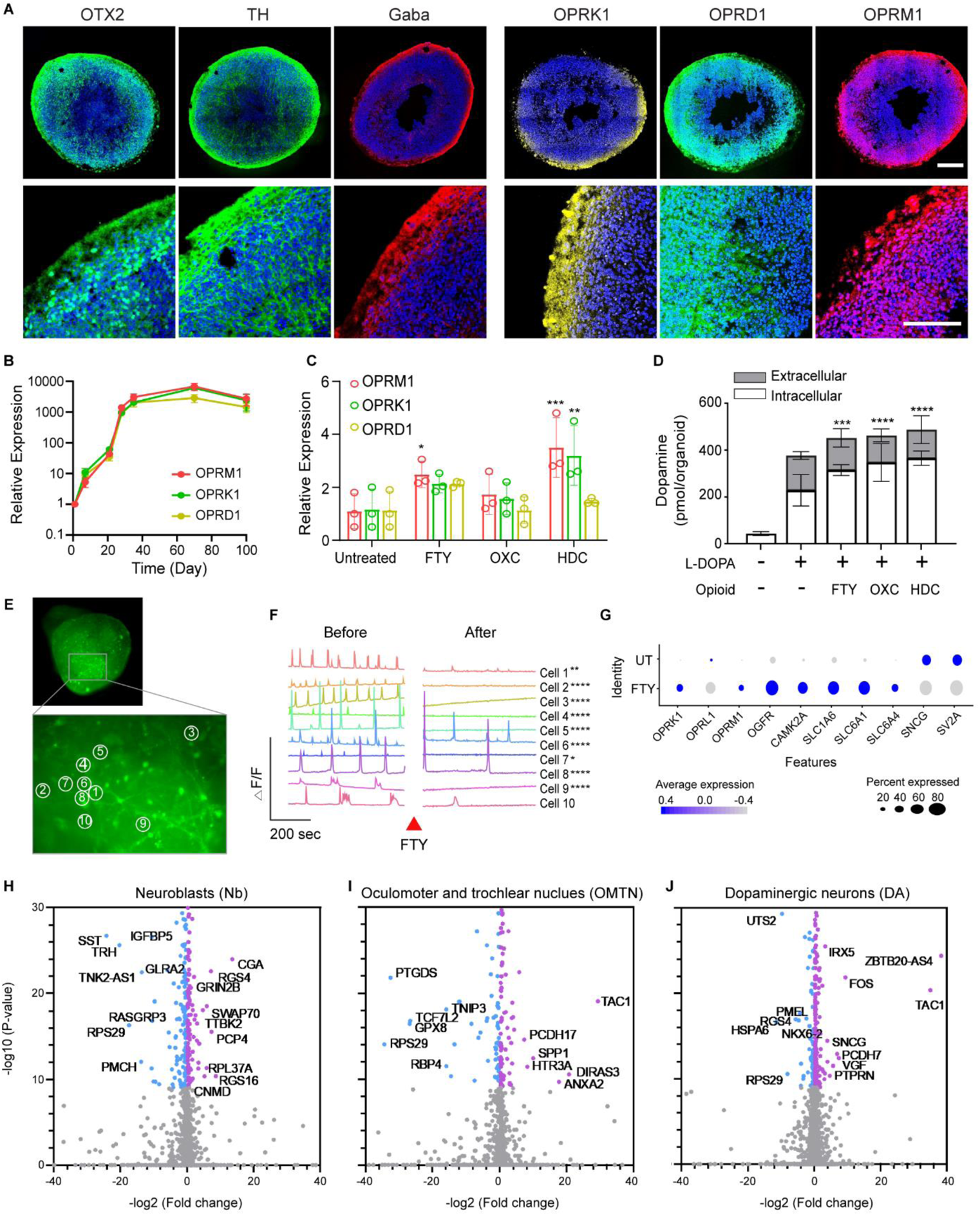
Cellular responses to acute opioid exposure for 4 hours. (A) Gene expressions of opioid receptors including mu (OPRM1), kappa (ORPK1), and delta (OPRD1) for 90 days of culture (n=5-7 organoids from 3 independent experiments). (B) ICC staining of opioid receptors of Day 35 organoids. (C-J) Results of 4-hour treatment of opioids (FTY; fentanyl, OXC; oxycodone, HDC; hydrocodone) in midbrain-like organoids at day 90. (C) Alterations in gene expression level of opioid receptors (n=3 organoids) and (D) dopamine synthesis and release in response to 4-hour opioids treatment (n=3 organoids). Statistical analysis was determined by One-way AVOVA as compared to untreated control (*p<0.05, **p<0.01, ***p<0.001, and ****p<0.0001). (E and F) Calcium imaging using Fluo4-AM of Day 180 organoids showing decreased frequency of calcium transients after fentanyl treatment (Movie S1). Statistical analysis of frequency and amplitudes was determined by t-test (*p<0.05, **p<0.01, and ****p<0.0001). Frequencies of all cells were significantly decreased after the FTY treatment (***p<0.001). (G) Summarized changes in the expression of opioid receptors (OPRK1, OPRL1, OPRM1, OGFR), sec ond messenger effects of Ca2+(CAMK2A), neurotransmitter transporters (SLC1A6, SLC6A1, SLC6A4), a nd synaptic plasticity (SNCG, SV2A). Each dot color and size represent relative gene expression level and proportion of expressing cells, correspondingly. (H-J) Volcano plots of gene expression changes in three cell types after fentanyl treatment. Blue: UT, Purple: FTY.

Acute opioid administration occurs in many clinical situations, including as pain relief during labor and delivery. To investigate the impact of short-term opioid treatment on the fetal brain (i.e., acute opioid exposure *in utero*), the midbrain organoids were treated with synthetic opioids (fentanyl, oxycodone, and hydrocodone) for 4 hours (**Figure S4**). The 4-hour treatment of opioids up-regulated the bulk mRNA expression levels of all three opioid receptors by 2 to 4-fold as compared to untreated organoids (**Figure 2C**). In addition, the intracellular and extracellular levels of dopamine also increased in organoids treated for 4 hours with fentanyl and for a subsequent hour with L-DOPA (**Figure 2D**). Measurements with Fluo-4 AM showed that the frequency and amplitude of spontaneous calcium transients were attenuated in cells in fentanyl-treated organoids (**Figure 2E, 2F** and **Movie S1**), consistent with the mechanism underlying opioids’ modulation of signal transmission and pain perception^53^.

Next, we performed single cell transcriptome analysis of fentanyl-treated organoids. We observed that the 4-hour fentanyl treatment altered gene profiles of neuroblasts (Nb), oculomotor and trochlear nucleus (OMTN), and dopaminergic neurons (DA) (**Figure 2 H-J**, **Figure S3**), but not in other neuronal types, such as Gaba, Sert, and red nucles (RN). Interestingly, when we examined the single cell expression of previously reported opioid response genes^27, 54^, synaptic plasticity genes (SNCG, SV2A) were expressed at a lower level in the FTY group (**Figure 2G**). This indicates that synapse transmission was downregulated by acute opioid treatment. Gene set enrichment analysis, however, showed no significant change in biological pathways, possibly because we sequenced the organoids on the same day of the acute treatment and ground level changes in gene expression profiles were not yet established. Overall, our results suggest that even a short-term opioid treatment elicits cell-type specific responses in the midbrain, but does not significantly alter the gene profiles in neuronal lineage specification and neuron projection.

### Chronic fentanyl treatment impairs neurodevelopment

Several neuroimaging and animal studies have shown the adverse effects of prenatal opioid exposure on neurodevelopment^26, 55^, including dysregulated functional connectivity and decreases in brain volume across multiple regions such as the midbrain, and many others have linked this exposure to clinical observations of delayed neurodevelopment ^19, 20, 56^. Therefore, we sought to investigate the effects of chronic fentanyl treatment using our midbrain organoid model, in which organoids were treated with 74 nM fentanyl for 53-77days starting from the first day of floor plate induction (Figure 1A). We first examined the morphology of the midbrain organoid over its development. Our time course analysis of midbrain organoids indicated that chronic fentanyl exposure did not change the gross morphology nor the size of the organoids during 77 days of culture (**Figure 3A**), possibly because organoid growth is eventually limited to a certain size anyway without the support of a 3D hydrogel scaffold and perfusable vasculature. Furthermore, there were no significant changes in the distribution pattern of OTX2 and TH proteins in the midbrain organoids, forming rosette-like regions (**Figure 3B**).

**Figure 3.**
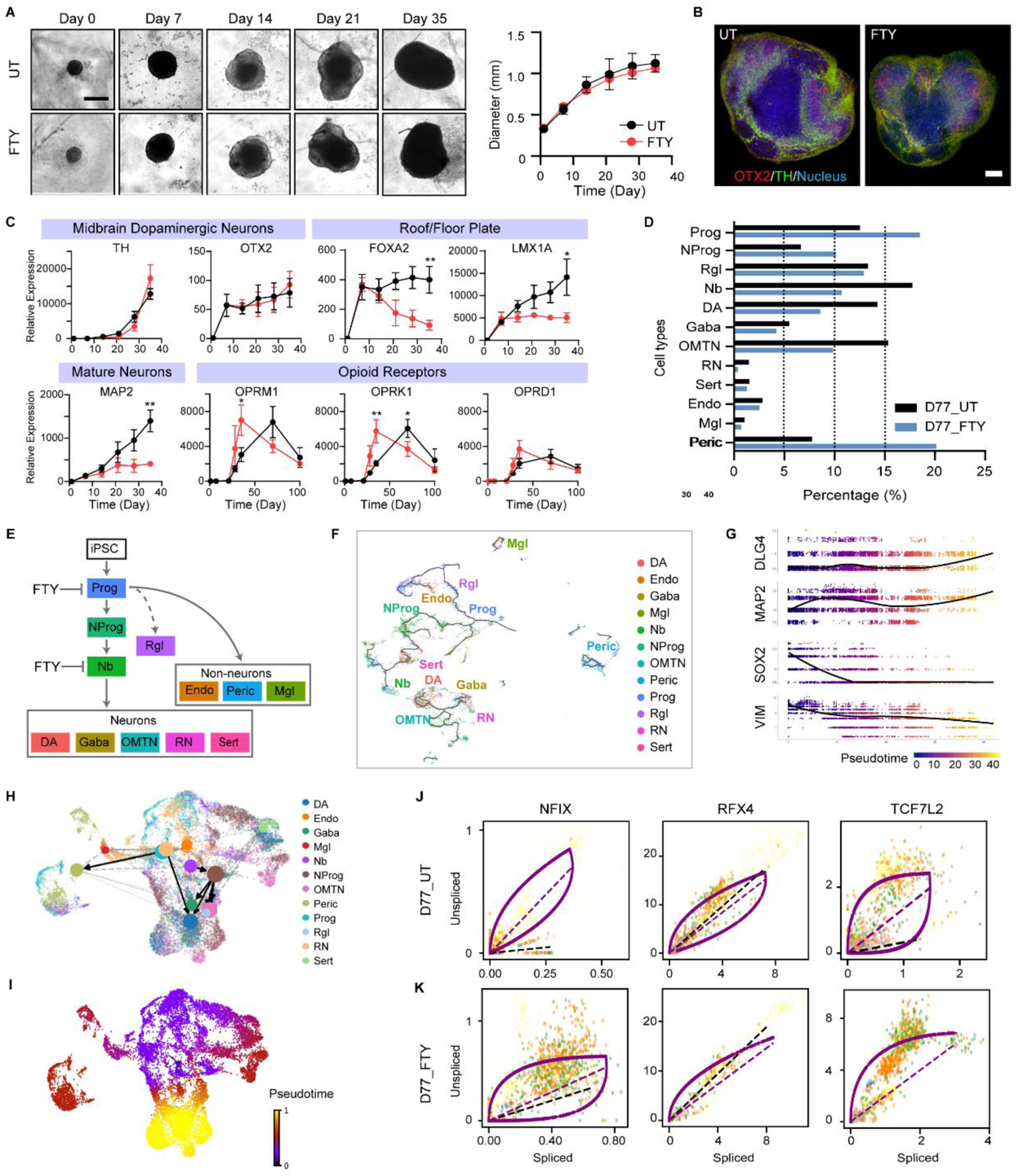
Chronic Fentanyl Treatment Altered Cell Heterogeneity in the Developing Human Midbrain Organoids. (A) Timeline of specimens showing organoid size and morphology (scale bar, 500 μm). Profiles of organoid growth (n=20 organoids, three individual experiments). (B) Spatial characterization of the midbrain patterning gene (OTX2) and function gene (TH) in Day 21 midbrain organoids (scale bar, 250 μm). (C) Fold changes of target genes of roof/floor plate (FOXA2, LMX1A), dopaminergic neurons (TH, OTX2), mature neurons (MAP2), and opioid receptors (OPRM1, OPRK1, OPRD1) (n=5-7 organoids per group/batch, three individual experiments) (t-test, *p<0.05 and **p<0.01). (D) Bar chart depicting the percentage of cells in untreated or fentanyl-treated midbrain organoids at Day 77. (E) Hypothesized hierarchy of neural cell development. Fentanyl affected cell fate transition in progenitors and neuroblasts. (F) Single cell lineage trajectory as plotted by similarity-based analysis (Monocle3 algorithm). (G) Trend of mature neuron markers (DLG4, MAP2) and stem cell markers (SOX2, VIM) of single cells in pseudotime (Monocle3 algorithm). (H) Predicted cell fate transition by RNA velocity (scVelo algorithm). (I) Predicted pseudotime state of single cells by RNA velocity. (J-K) Phase portrait showing the levels of unspliced and spliced mRNA in untreated or fentanyl treated organoids. Above or below the purple dashed line indicates increasing or decreasing expression of a gene.

Next, we explored whether the differentiation and maturation of various cell types in the midbrain organoids were affected by chronic opioid exposure. Interestingly, the expression level of both midbrain progenitor markers, FOXA2 and LMX1A, and pan-neuronal markers, MAP2, was largely lower in the chronic fentanyl treatment group than in the untreated organoids (**Figure 3C**). In contrast, there was no significant difference in the expression of other genes related to pluripotency (OCT4, NANOG), midbrain (DAT, PITX3), forebrain (PAX6), or hindbrain (TBR2, GBX2) regional identity at the bulk mRNA level, as measured by RT-qPCR (**Figure S6A**). All three opioid receptors in the chronic fentanyl exposure group were expressed much earlier than in untreated organoids (**Figure 3C**). The expression of the opioid receptors in the chronic fentanyl treatment group peaked at day 35 while untreated organoids showed the highest expression levels of opioid receptors at day 70.

We next examined scRNA-seq data to further characterize how fentanyl affected midbrain neuronal differentiation. Comparing the cell type composition, a lower percentage of mature neurons (DA, Gaba, OMTN, RN, Sert) and a higher percentage of pericytes (Peric) and progenitors (Prog, NProg) were observed in the fentanyl-treated samples (D77_FTY) as compared with the untreated samples (D77_UT) (**Figure 3D**, **S2A**). Further, mature dopaminergic neurons (DA) accounted for 14.26% of cells in the D77_UT sample and only 8.58% of cells in the D77_FTY sample. However, although less mature neurons developed in the fentanyl-treated samples, the average level of TH remained similar.

We hypothesized that the change in cell composition resulted from alterations in cell lineage decisions in midbrain development (**Figure 3E**). To understand the chronological neurodevelopment, we constructed the developmental cell fate trajectories by two state-of-art lineage decision methods: Monocle 3 (similarity-based method, **Figure 3F-G**) and scVelo (RNA velocity-based method^48, 57^, **Figure 3H-J**, **Methods**). In Monocle 3, UMAP^58^ visualization of our overall dataset showed two distinct groups: progenitors (Prog, NProg, Rgl, Nb) and neurons (DA, Gaba, OMTN, RN) (**Figure 3F and Figure S7 A-C**). Pseudotime lineage analysis showed the dynamic gene profile changes over pseudo-age. For instance, stem cell genes (SOX2 and VIM) and mature neuron markers (MAP2 and DLG4) had an inverse relationship, changing in opposite directions (**Figure 3G** and **Figure S7 D-F**). Furthermore, we calculated RNA dynamics (i.e. rates of transcription, splicing, and degradation of individual genes) based on the mapped spliced and unspliced mRNA reads with scVelo (**Figure S8A-B, Methods**). By quantifying the connectivity of cell clusters, partition-based graph abstraction (PAGA) provided a simple abstract graph of cell fate connectivity (**Figure 3H**). We double checked the pseudotime projection of our dataset and plotted out velocity streamlines of RNAs in both a normal brain organoid (**Figure S9A**) and a growth-impaired organoid (**Figure S9B**). In theory, quiescent stem cells and terminally differentiated cells (e.g. neurons) show close to zero velocity values as they are at a steady state of spliced and unspliced mRNA levels. The size of the stream line arrows was in proportion to the value of the cell’s velocity. We observed low velocity fields in a subpopulation of Prog and Rgl, as well as in Gaba and DA (**Figure S9A-B**), which is consistent with the findings of previous studies^48, 59^. In addition, phase portraits of neuronal specification markers and mature neuron function markers showed that chronic fentanyl exposure arrested cells at an early stage of neurodevelopment (**Figure 3J-K**, **S9C-D**). Although radial glia-like cells were identified in midbrain organoids, their specification and activity to give rise to certain neurons are yet to be clarified.

Overall, our trajectory maps illustrated that: 1) neuron specification and maturation was affected by the chronic fentanyl exposure, and more cells arrested as Prog or Nb instead of becoming neurons (**Figure 3D-E**); 2) distinct neuron lineages converge back to intermediate progenitors (i.e. neuroblasts); 3) iPSCs give rise to both mesoderm-derived cells and ectoderm-derived neural cells in midbrain organoids. Together, these data suggest our midbrain organoid model as a comprehensive framework to investigate cell lineage decisions and driver gene dynamics in neurodevelopment.

### Identification of gene regulatory network for major cell types

To find the master regulators that control cell fate decisions, we performed regulatory network inference and clustering on scRNA data using SCENIC R package^49^ (**Methods**). Instead of looking at individual genes in the differentially expressed matrix, this method examines gene networks, taking cis-regulatory motifs and transcription factors into account. In brief, we scored the activity of each regulon in all single cells and identified the top transcription factor networks (**Figure 4A**, **Figure S10**). Both untreated and fentanyl-treated samples showed a significant enrichment in CREB5 (25 predicted target genes) and MAX (10 predicted target genes) regulons (**Figure 4B**, **Figure S5**). CREB5 is a cyclic-AMP response element binding (CREB) protein that has roles in neuronal survival, differentiation, memory, and drug addiction^60^. MAX regulates more than 180 genes in the human telencephalon and contributes to the development of all regions in the brain^61^. Interestingly, progenitor cells (Prog, NProg, Rgl) in the treated sample (D77_FTY) showed lower levels of regulon activity (SOX2, BARHL2, FOXJ1, RFX4, ETV1) for neuronal differentiation, specification, migration, and survival (**Figure 4A, 4C-D**, **Figure S10C**). BARHL2 contributes to subtype specification and neuron migration^62^, while ETV1 promotes dendrite and synapse formation in granule neurons^63^. A broad spectrum of transcription factors for neuronal function genes was identified in the untreated sample, such as OTP (dopaminergic neuron development), ISL1 (required for motor neuron identity and motor axon guidance), HDAC6 (synaptic plasticity), and ZMIZ1 (neuron maturation) (**Figure 4A, 4G**, **Figure S10C**). We do not observe significant differences in the regulon activity of neurons or mesoderm-derived cells between the treated and untreated samples (**Figure S10 A-B**). In contrast, progenitor cells in the fentanyl-treated sample (D77_FTY) showed higher activities in regulons (SOX9, NFIX, TCF4) for early neurodevelopment patterning and differentiation (**Figure 4A**, **Figure S10C**), suggesting those progenitors were at an early stage of neurodevelopment. HMGN3 (stress-responsive), DDIT3 (pro-apoptotic), and ZNF91 (apoptosis during early neuronal differentiation) regulons were also found to be active in the treated sample, implying that the cells are under stress. Our results traced the gene regulatory networks that were causing the cell fate divergence in progenitor cells (Prog, NProg, Rgl) and identified potential regulators for each single cell.

**Figure 4.**
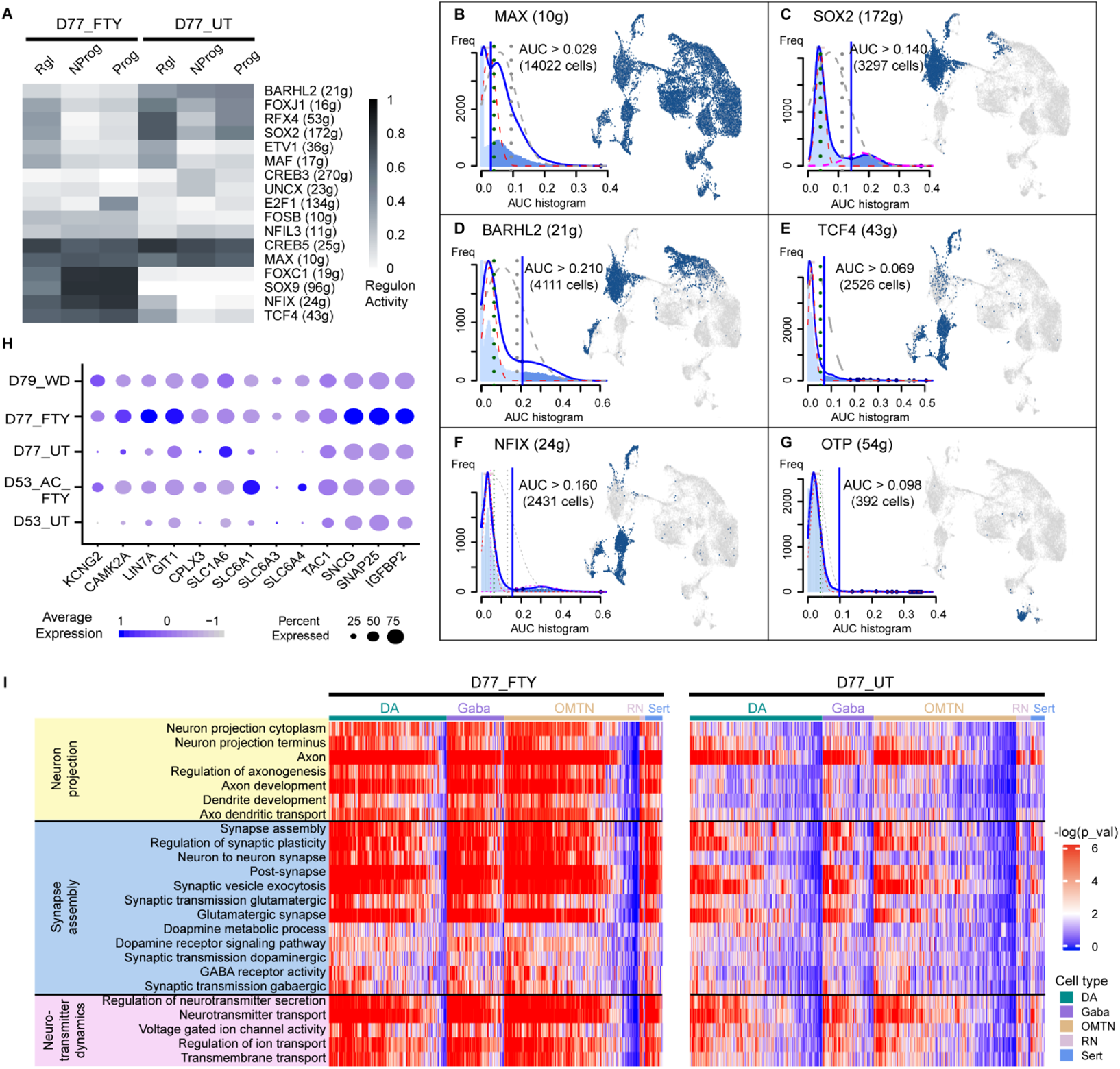
Chronic fentanyl exposure alters regulon networks and synaptic activity. (A) Identified master regulators in each cell type based on regulon analysis (SCENIC algorithm). The number of predicted target genes was given for each transcription factor in the bracket. (B-G) Activity score of each regulon network and its binarized activity mapped in UMAP. The blue vertical line in the AUC histogram yielded the cutoff for the “ON” or “OFF” state of that regulon network. Both samples (D77_UT and D77_FTY) showed high activity of the MAX regulon. Untreated sample (D77_UT) showed a higher percentage of cells with “ON” states of SOX2, BARHL2, and OTP, while fentanyl-treated sample (D77_FTY) had higher activities of early neurodevelopmental transcription, such as TCF4 and NFIX. (H) The dotplot showed the individual gene expression level and percentage of expressing cells in the neurons of all five samples. A higher percentage of cells in D77_FTY and D53_AC_FTY were expressing genes related to ion transport (KCNG2), synaptic plasticity (CAMK2A, LIN7A, GIT1, CPLX3), and neurotransmitter release (SLC1A6, SLC6A1, SLC6A3, SLC6A4) as compared to untreated samples. (I) Single sample gene set enrichment analysis (ssGSEA) showed altered pathways relating to neuron projection (yellow), synapse assembly and signal transduction (blue), and neurotransmitter dynamics (purple) in fentanyl-treated samples (left heatmap) and untreated samples (right heatmap).

### Chronic fentanyl exposure changes neuronal communication in synaptic activity

We analyzed the heterogeneous cell type-specific response to chronic fentanyl exposure based on single cell RNA sequencing results. Differentially expressed genes between D77_UT and D77_FTY were identified by using the “FindMarkers” function in Seurat. Both samples expressed high levels of VGF, SCG2, VAMP2, and RAB3A in various neuronal types (DA, Gaba, OMTN, Sert, RN) (**Figure S11, S12**), which are secretory- or vesicle-associated proteins that regulate neurotransmitter release^64, 65^. A higher percentage of neurons (DA, Gaba, OMTN, Sert, RN) in D77_FTY expressed genes related to neuronal functions such as ion transport (KCNG2), synaptic plasticity (CAMK2A, LIN7A, GIT1, CPLX3), and neurotransmitter release (SLC1A6, SLC6A1, SLC6A3, SLC6A4) (**Figure 4H**). Interestingly, opioid receptor genes (OPRD1, OPRK1, OPKL1, OPRM1) did not present significantly different expression at a single cell level in the chronic fentanyl sample. Our ssGSEA analysis also identified pathways with gene set enrichment in each single cell (**Figure 4I**, **Methods**). In the D77_FTY sample, gene sets related to synaptic activity, neuron projection, and neurotransmitter transport were highly enriched in neurons (DA, Gaba, OMTN). Differential responses between DA and Gaba were not observed (**Figure 4I**, **S11**), likely because neural circuits were not yet well established in organoids. In addition, progenitors (Rgl, Prog, NProg, Nb) in D77_UT exhibited high regulon activity in neuron specification and migration, while progenitors in D77_FTY showed high activity in neurogenesis, differentiation, and axongenesis (**Figure S10B**). This indicates that progenitors in the D77_FTY arrested at a much earlier stage than those in D77_UT.

Mesoderm-derived cells (pericytes, endothelial cells, and microglias) in untreated midbrain organoids (D77_UT) were enriched with genes that mediate endothelium development, angiogenesis, and extracellular structure organization. In chronic fentanyl-treated organoids, we observed an elevated expression in genes related to immune response (PLCG2, TFF3, CRYAB) and homeostasis (EPYC, SPINK6) (**Figure S13**). Interestingly, pericytes were highly responsive to fentanyl treatment, which is demonstrated by the significant transcriptome changes as compared to untreated pericytes (**Figure S3A, S12A**).

Furthermore, the extracellular dopamine level of organoids with the chronic fentanyl treatment was higher than that of untreated organoids (**Figure S6B**), but not significantly. The organoids with chronic fentanyl treatment also showed less neurons with calcium signaling (**Figure S6C** and **Movie S2**), indicating that cellular excitability and signal transmission were affected by fentanyl.

### Neural development resumes in transcriptomes when fentanyl is withdrawn

To determine if and to what degree the impact of fentanyl exposure on neurodevelopmental program can be reversed when fentanyl is withdrawal after chronic exposure, we alleviated organoid stress by withdrawal of fentanyl for 2 days. We performed similar differential expression analysis between the untreated sample (D77_UT) and withdrawal sample (D79_WD), and found that regulon activities (FOXJ1, RFX4, SOX2, BARHL2) of neuronal differentiation and subtype specification were substantially restored in progenitor cells (Prog, NProg, Rgl) (**Figure 5A-B**). We also observed an increased expression of ventral midbrain progenitor genes (LMX1A), subtype specification genes (NKX6-1, FOXA2), and mature neuron markers (RBFOX3, NFE2L1, HMGN3) in the D79_WD sample as compared to D77_FTY (**Figure 5C**). Gene set enrichment analysis showed that the synaptic activity and neuron projection in D79_WD were more similar to D77_UT than D77_FTY (**Figure 5D**), indicating that the synaptic activity decreased to a basal state in response to drug withdrawal. All the above data suggest that neural development resumes in transcriptomes when opioids are withdrawn.

**Figure 5.**
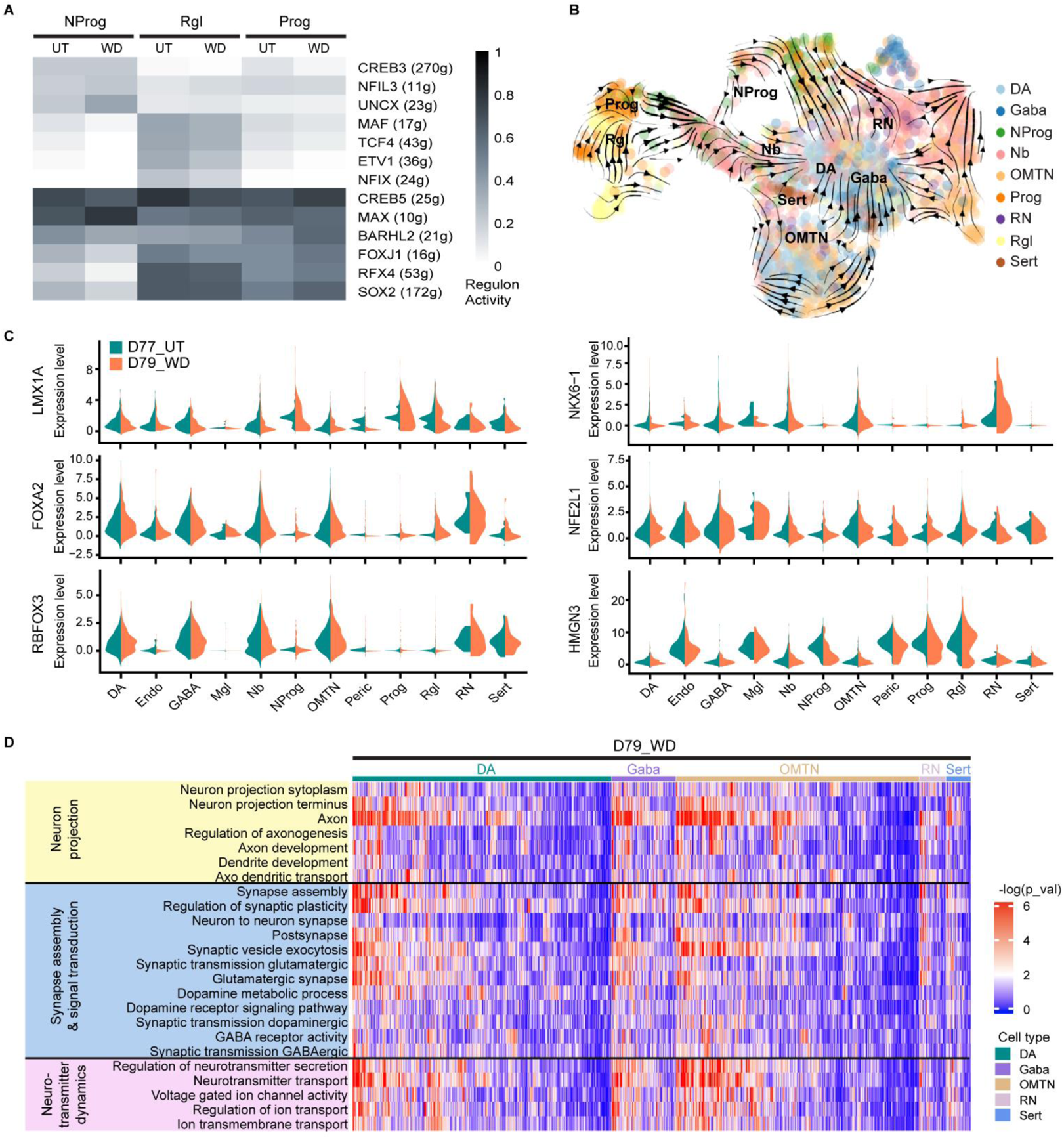
Neural development resumes in transcriptomes when fentanyl is withdrawn. (A) Key master regulators identified in withdrawal sample (D79_WD) versus untreated sample (D77_UT) (SCENIC algorithm). (B) RNA velocities were visualized as streamlines in a UMAP-based embedding for all five samples. Spliced/unspliced mRNA dynamics disentangled cell lineage commitment at the single cell level. (C) The violin plots show the gene expression level and percentage of expressing cells in D77_UT and D79_WD samples. (D) Single sample gene set enrichment analysis (ssGSEA) showed pathways relating to neuron projection (yellow), synapse assembly and signal transduction (blue), and neurotransmitter dynamics (purple) restore basal states in fentanyl withdrawal samples. The pattern was more similar to untreated samples (D77_UT, right heatmap of Fig. 4I) than fentanyl-treated samples (D77_FTY).

## Discussion

In utero opioid exposure has been reported to impair neurodevelopment and potentially cause poor neurocognitive, behavioral, and developmental outcomes in childhood^19, 20^. However, the mechanism underlying this dysregulation is unclear, and more focused research must be done. One of the primary adult neural circuits involved in opioid action is the mesolimbic reward system. This system generates dopamine signals that arise in the ventral tegmental area (VTA) in the midbrain and propagate throughout the rest of the brain, playing a role in learning and reward pathways. Drugs of abuse, including opioids, take advantage of this system, inducing dopamine surges and relieving pain through acute opioid exposure^66^. Chronic opioid exposure, however, does not simply have an exaggerated effect – it disrupts dopamine signaling and causes transcriptional and epigenetic changes in the brain regions within the mesolimbic system, promoting addiction and vulnerability to relapse^66, 67^. Although extensive clinical and preclinical data have demonstrated that opioid addiction is strongly associated with the mesolimbic system in an adult brain, little is known about the particular association in the developing human fetal brain. We therefore aimed to use midbrain organoids to investigate the molecular and cellular alterations caused by chronic opioid exposure in the midbrain region, which is the largest dopamine-producing area in the brain and heavily involved in the mesolimbic reward system, during neurodevelopment.

Our midbrain region-specific organoids were generated by a guided method using midbrain patterning molecules. Dual-SMAD inhibition factors, LDN 193189 and SB431542, and a Wnt pathway activator, CHIR 99021, instructed iPSC differentiation toward floor plate lineages. Subsequent treatment of a small molecule agonist, purmorphamine, and recombinant SHH boosted floor plate cells toward ventral mesencephalic lineages. We compared the gene expression profiles of organoids in multiple independent batches (Figure S1) that were generated using two different iPSC lines (i.e. FA11 and QR19). No significant variations were found, suggesting that our guided method could generate reliable and consistent midbrain organoids. In addition, our method does not involve embedding the organoids in matrices (e.g. Matrigel) for high reliability and reproducibility of organoid generation^41, 68^.

These midbrain organoids also exhibited the cytoarchitecture and compositional features unique to the developing human ventral midbrain region in vivo^52, 69^. The electrophysiological properties of the organoids at day 35 indicate that neurons located at the surface of the organoids are not fully mature, as evidenced by their depolarized resting membrane potential, slow or absent pacemaking activity, and inability to maintain long action potential trains compared to mature neurons in an adult human brain^41, 70, 71^. This is not surprising, however, as organoids model early stages of human embryonic development, in which fully realized electrophysiological properties, such as action potentials and spontaneous synaptic transmission, are not yet observed and ontogenetic changes still occur^72^. Indeed, electrophysiological properties of neurons in midbrain organoids can gradually become more mature^71^, and cortical organoids have been shown to parallel the timeline of neurodevelopment in vivo, using genome-wide analysis of the epigenetic clock and transcriptomics to demonstrate that organoids reached postnatal stages around day 300 of culture in vitro^68^. We believe that midbrain organoids could develop and mature over time to similarly reflect in vivo neurodevelopment.

Once midbrain organoids were established as viable models to investigate midbrain neurodevelopment, fentanyl, a potent synthetic opioid, was used to treat the organoids. Acute fentanyl treatment (74 nM for 4 hours), increased opioid receptor expression and dopamine release, but did not have time to elicit significant changes in gene profiles related to cell lineage development and neuron projection. Chronic fentanyl treatment (74 nM for 55-77 days), however, did affect the cell fate of iPSCs. Leveraging single-cell transcriptome profiling to identify molecular responses at the cellular level, we showed that chronic fentanyl exposure impaired the neurodevelopment of the midbrain organoids, as demonstrated by an increased neuroprogenitor pool and delayed neuronal differentiation and maturation. This neurodevelopmental dysregulation can possibly lead to disrupted brain regionalization and poor cognitive function. Moreover, a series of genes and pathways related to neural development showed significantly altered regulation in long-term fentanyl-treated organoids.

Our findings are consistent with the clinical observation of impaired neurodevelopment in neonates with in utero chronic opioid exposure. The delayed neurodevelopment might be associated with clinical observations including delays in psychomotor development, lower IQ scores, and higher total behavioral problem scores in the childhood^19, 20^. In addition, neonates often suffer from unusual withdrawal symptoms including high-pitched crying, poor feeding, exaggerated Moro reflex, irritability, and trouble sleeping^73^. These symptoms, caused by neonatal abstinence syndrome (NAS), are exacerbated by a sudden discontinuation of opioids upon birth. Although our results indicate that synaptic activity was decreased to a basal state and neurodevelopment resumes in transcriptomes at fentanyl withdrawal, NAS symptoms persist by affecting both the central and autonomic nervous systems, as well as the gastrointestinal system^74^. These findings can hopefully lead to future, viable therapeutics for both in utero chronic opioid exposure and NAS.

## Limitations of Study

Challenges remain in uncovering native brain functionality and correlating organoid-based findings to clinical readouts. Primarily, brain organoids still lack the full cellular maturation, heterogeneity, architecture, vasculature, and microenvironmental niche seen in the actual human brain.^44, 75, 76^ This limits the extent to which organoids can be a fully realistic model and demands that organoids undergo further characterization to better understand these limitations. Recent advances in spatial multi-omics, along with high-throughput transcriptomic and proteomic profiling, offer great opportunities to characterize the developmental and functional dynamics of brain neurogenesis and synaptogenesis in organoids.^77–79^ Future studies could also validate these models through human studies or animal models.

Another challenge is the scope of region-specific organoids. Neuropsychiatric disorders, including substance abuse, affect several brain regions and implicate complex neural circuits.^80–82^ Midbrain organoids, although providing valuable information regarding the effects on the midbrain, lack the functional neural circuits found in the native central nervous system (CNS) and can inherently only tell part of the story. Until further advancements in brain organoid development, systematic studies using several different region-specific organoids are important to fill out the picture.

A final challenge is that brain organoids can only reveal the effects of a single drug abuse and identify those potential biomarkers and therapeutic targets. In the event of polydrug abuse, these models are unable to match between specific downstream effects and their drug of origin. Multiples studies must be done in parallel, each investigating an individual drug abuse, to appreciably gain a mechanistic understanding of each drug.

Overall, long-term exogenous opioid treatment elicits massive transcriptomic profile changes in neurodevelopment and neuronal communication. Future studies in correlating the clinical readouts with transcriptomic markers would establish comprehensive evaluation of drug abuse outcomes and offer opportunities in therapeutic targets.

## Conclusion

In midbrain organoids, a short fentanyl treatment induced an increase in dopamine release without significant changes in the gene expressions of cell lineage development. In contrast, chronic fentanyl exposure impaired cell subtype specification and altered synaptic activities in neurons, indicating arrested neurodevelopment. Upon fentanyl withdrawal, the neurodevelopment resumed to normal at the transcriptomic level. Overall, our study dissected the molecular landscape of opioid responses in neural and non-neural cells and unveiled the affected pathways that may inform strategies to boost treatments in substance abuse and neonatal abstinence syndrome.

## Methods

### Culture of human induced pluripotent stem cells (iPSCs)

The FA11 iPSC line was generously provided by the Columbia Stem Cell Initiative with an approved IRB. FA11 cells were derived from human dermal fibroblasts of a healthy donor ^83^. The iPSCs were maintained under feeder-free conditions over Matrigel-coated plates in mTeSR Plus media (Stemcell Technologies). The media was changed daily and the iPSC cultures were split into 1:6-1:10 every 5 days using ReLeSR (Stemcell Technologies).

### Generation of midbrain organoids

iPSCs before passage 30 were used to generate the organoids. To form embryoid bodies (EBs), iPSCs were dissociated into single cells with accutase (Gibco) treatment and 8,000 cells were plated in each well of non-treated U-bottom 96-well culture plates (Corning) with mTeSR plus media and 10μM ROCK inhibitor Y27632 (Tocris Bioscience). In brief, to induce iPSC differentiation toward a floor plate, the EBs were treated with dual-SMAD inhibition factors, LDN 193189 and SB431542, and a Wnt pathway activator, CHIR99021^84, 85^. For efficient midbrain patterning toward a ventral mesencepthalic fate, Sonic Hedgehog (SHH) signaling was subsequently activated by a small molecule agonist, purmorphamine, in combination with recombinant SHH^86^ (**Figure 1A**). Detailed product information could be found in **Table S2**.

On day 0, we used neuronal induction medium, which composed of 15% Knockout serum replacement (Gibco), 1% GlutaMax (Gibco), 1% minimum essential media-nonessential amino acids (MEM-NEAA) (Gibco), and 0.1% β-mercaptoethanol (Gibco) in Knockout DMEM/F12 (Gibco), supplemented with 100nM LDN193189 (Stemgent) and 10μM SB431542 (Tocris Bioscience). On day 2, we changed to the neuronal induction media that was supplemented with LDN193189, SB431542, 100ng/ml SHH (R&D Systems) and 2μM Purmorphamine (Calbiochem). On day 4, 3μM CHIR99021 (Stemgent) was added to the medium. From day 6 to day 8, the basal media containing LDN193189 and CHIR99021 was gradually replaced to a differentiation media containing DMEM/F12: Neurobasal (1:1) (Gibco), N2 supplement (Gibco), B27 supplement without vitamin A (Gibco), 1% GlutaMAX, 1% MEM-NEAA, 0.1% β-mercaptoethanol and 1% penicillin-streptomycin (Gibco). On day 9, the differentiation media was supplemented with CHIR99021, 10ng/ml BDNF (Prospec), 0.2mM ascorbic acid (Sigma), 20ng/ml GDNF (Prospec), 0.2mM dibutyryl-cAMP (Calbiochem), 1ng/ml TGF-β3 (R&D Systems), and 10μM DAPT (Tocris). From day 11, the organoids were cultured in the same media without CHIR99021. Media was then changed every 3-4 days. On day 21, the organoids were transferred into non-treated 24-well plates by using a 200μl tip with a wide bore opening. The organoid growth was monitored by bright-field microscopy (Nikon Eclipse) for 55 days, and the diameter was measured using Image J.

### Acute and chronic opioid treatment

Day 95-organoids were treated with fentanyl (FTY, Cerilliant, F-002), oxycodone (OXC, Cerilliant, O-002), and hydrocodone (HDC, Cerilliant, H-003) at a concentration of 4 nM, 7 nM, 19 nm, 37 nM and 74 nM, respectively, for 4 hours (“acute” opioid treatment). The cytotoxicity was assessed by CellTiter-Glo^®^ Luminescent Cell Viability Assay (Promega). The changes in the mRNA expression of opioid receptors and the dopamine release levels caused by opioid treatment were evaluated by real-time PCR and HPLC analysis, respectively, as described below. Fentanyl (74 nM in culture media) was also used to treat organoids starting on day 1 of the neuronal induction (“chronic” opioid treatment). On day 77, to analyze the effects of opioid withdrawal, media was replaced to fresh media without fentanyl, and the organoids were incubated for two days prior to single cell RNA sequencing analysis. (**Table S1**)

### Immunohistochemistry

The organoids were fixed in 4% paraformaldehyde for 1 hour, saved in a 30% sucrose solution in PBS overnight, and subsequently embedded in O.C.T. compound (Tissue-Tek) for cryosectioning. Frozen organoids were cryosectioned at a thickness of 20μm. We heated the slides up to 95 °C in a citrate buffer (10mM Sodium Citrate, 0.05% Tween 20, pH 6.0) for antigen retrieval. For immunohistochemistry, the slides of organoid cryosections were permeabilized with 0.2% TritonX-100 in PBS 20min at room temperature, and then blocked with 3% Bovine Serum Albumin (BSA, Sigma) with 0.1% Triton X-100 in DPBS for 1 hour at room temperature. The sections were incubated with primary antibodies overnight at 4 °C and with secondary antibodies for 1 hour at room temperature (**Table S2**). All sections were stained with DAPI (Sigma) for cell nuclei. Images were taken on a confocal microscope (LSM 710, Zeiss).

### Real-time qPCR (RT-qPCR)

Gene expression profiles of the organoids during development were evaluated for 35 days by real-time (RT) PCR. Total RNAs were isolated from organoids using TRIzol^®^ reagent (Invitrogen) and reverse-transcribed using iScript^TM^ cDNA synthesis kit (Bio-Rad) to produce cDNA. Quantitative RT-PCR was performed using PowerUp^TM^ SYBR^TM^ Green Master mix (ThermoFisher) and QuantStudio^TM^ 3 Real-Time PCR System (Applied Biosystem). ΔΔCt method was applied to normalize expression levels of each gene to that of GAPDH. The sequences of primers were described in **Table S3**.

### Electrophysiology

Whole-cell patch clamp recordings were taken from organoids as previously described^87–89^. Day 35 organoids were attached to Matrigel-coated cover glasses overnight. Recordings were performed at 22°C under continuous perfusion (1 mL/min) with a bath solution containing: 119 mM NaCl, 5mM KCl, 30 mM HEPES, 2 mM MgCl_2_, 2 mM CaCl_2_ and 10 mM glucose. The solution was adjusted to 310 mOsm with sucrose and to pH 7.3 with KOH. Borosilicate glass pipettes with a tip resistance of 3–4 MΩ (G150F-4, Warner Instruments) were pulled on a P-97 Flaming-Brown micropipette puller (Sutter Instruments) and filled with an internal solution containing: 130 mM K-gluconate, 10 mM KCl, 2 mM Mg-ATP, 0.2 mM Li-GTP, 0.6 mM CaCl_2_, 5 mM MgCl_2_, 0.6 mM ethylene glycol-bis(2-aminoethylether)-N,N,N′,N′-tetraacetic acid (EGTA), and 5 mM HEPES titrated to a pH of 7.1 and an osmolality of 310. Neurons were visualized under a 40X water immersion objective by DIC optics (Olympus). Recordings were performed with an Axopatch 700B amplifier (Molecular Devices) and digitized at 10 kHz with an ITC-18 (HEKA Instruments). Data analysis was performed using Clampfit 10 software (Molecular Devices, Sunnyvale, CA) and Matlab 8.0 (MathWorks, Natick, MA). In each neuron, input resistance, resting membrane potential, and spontaneous firing frequencies were monitored throughout the recording, and only cells with neuronal morphology and a stable baseline activity that continued for >5 minutes were counted as tonically active. In order to determine the identity of recorded neurons, neurons were labeled with biocytin (Sigma-Aldrich) added to the patch pipette saline for >10 minutes, fixed in 4% PFA for 1 hour and immunostained with anti-TH antibodies.

### Calcium imaging

On day 180 of culture, both fentanyl-treated and untreated organoids were used for calcium imaging. The organoids were incubated with the cell-permeable calcium indicator Fluo-4 AM (ThermoFisher) for 30 min at 37°C. Time-lapse changes in Ca^2+^ levels in live organoids were imaged using a fluorescence microscope (Eclipse TS100, Nikon). To see the instant effect of the fentanyl treatment on the calcium channel, untreated organoids were recorded before and after the addition of fentanyl (74 nM in culture media). The organoids were imaged every 2 seconds at room temperature. The fluorescence intensity was determined by Image J and the data were normalized to ΔF/F_0_ using the following equation: y = (F_Δsec_ – F_0sec_)/F_0sec_.

### Dopamine release measurement

Intracellular and extracellular dopamine production from the midbrain organoids was characterized by high-performance liquid chromatography (HPLC). First, three organoids per condition were transferred to a 96-well plate on Day 55. L-Dopa (100 μM) was added to each well and incubated for 1-2 hours. For extracellular dopamine measurements, media was removed and washed with PBS, followed by the immediate addition of Tyrode’s containing KCl (40 mM). After 20-30 min of incubation, the sample from each well was collected and mixed with 0.12 M perchloric acid at a 1:1 (v/v) ratio. For intracellular dopamine measurements, all media was replaced with Tyrode’s containing KCl and organoids were then dissociated. The lysate was mixed with 0.12 M perchloric acid at a 1:1 (v/v) ratio. After 10-15 min incubation, cell debris was removed by centrifugation at 10,000 g and 4 °C, and the supernatant was collected for HPLC.

Organoid dopamine levels were then determined by HPLC with electrochemical detection, as previously described.^90, 91^ In brief, samples were separated on a VeloSep RP-18, 3 μM, 100×3.2mm column (PerkinElmer, Waltham, MA) with a Gilson 307 HPLC piston pump set to a flow rate of 0.7 mL/min and a mobile phase containing 45 mM NaH_2_PO_4_, 0.2 mM EDTA, 1.4 mM HSA, 5% methanol, and pH 3.2. Dopamine was detected on an ESA Coulochem II electrochemical detector at 350 mV oxidation potential. Data was then collected using Igor software, and dopamine concentration was calculated from areas under the HPLC peaks using calibration curves.

### Single cell RNA sequencing

Five independent samples of midbrain organoids (**Table S1**) were gently dissociated to single cells with papain enzymatic dissociation (Worthington Biochemical Corporation). After filtering cell clumps with a cell strainer (35um mesh), suspended single cells were encapsulated with barcoded beads in oil droplets by 10X Genomics Chromium technology, as per the manufacturer’s manual. The resulting single cell 3’-end libraries were sequenced on Illumina® NovaSeq™ 6000 Sequencing System (2×100bp pair-end) at the Single Cell Analysis Core of the Columbia Genome Center. 10X genomics’ Cellranger pipeline v3.1.0 with human reference transcriptome GRCh38 was used to process the data. Details of software and algorithms used were in **Table S5**.

### Cell type annotations

The Seurat package (V3.0) in R (V3.6.3) was used to normalize the expression matrix and identify differentially expressed genes. The Glmnet package (V4.0) was used to assign cell types based on cells’ likelihood to an annotated human fetal midbrain dataset^52, 92–95^. First, we transformed both the experimental and reference datasets’ gene counts separately using the “sctransform” function in Seurat. Secondly, we used Seurat’s CCA integration method to account for technical differences between both datasets, and co-clustered both datasets in UMAP^58^. Then, we trained a multinomial logistic regression classifier (Glmnet R package) on the reference’s integrated data excluding the “unknown” (“Unk”) cell type and predicted the cell type of both datasets. In non-Unk reference cells, the classifier’s accuracy was 99%. To validate the cell types assigned, we compared the gene signatures of each cell type between each dataset using highly correlated or anti-correlated genes to predict the cell types in our samples. Our results showed similar gene patterns compared to the reference dataset. Overall, we found that the cell type composition of the reference dataset and our own dataset were similar, with some variations in percentages of progenitor cells, neuroblasts, and oligodendrocyte progenitor cells (OPCs) (**Figure 1F**, **S2**). We grouped the sub-clusters of a same cell type into one major type. For instance, any cell type with hDA in the name was mapped to the DA major type. Any cell type with Prog in the name was mapped to the Prog major type. Any cell type with hNb in the name was mapped to the Nb major type. A detailed table of cell metadata table was deposited in Gene Expression Omnibus (GEO) under accession number GSExxxxxx (This number will be available following the acceptance of the manuscript).

### Differential expression analysis

Differentially expressed genes between samples or cell types were identified by using the “FindMarkers” function in Seurat. The following parameters were used: min.pct = 0.25, logfc.threshold = log(2). The top genes in each comparison group were selected with a pre-filtering step (p_val_adj < 1e-10, pct.1 > 0.1, pct.2 > 0.1, avg_logFC>5 or <-5) to remove low count and insignificant genes, and then ranked by the absolute value of log fold change. Heatmap of top differentially expressed genes were shown in Figure S3.

### Single sample gene set enrichment analysis (ssGSEA)

We performed ssGSEA to identify the activated pathways in our samples using the pipeline previously described in identifying subtypes with enriched gene sets^96^. In brief, normalized gene expression matrix (generated by Seurat sctransform function) and gene sets of interest (from GSEA Molecular Signatures Database v7.2 and Gene Ontology Database) were used as inputs. Enrichment scores of a certain gene set were calculated for both the experimental dataset and a random permutated dataset of 1000 cells. P-values of each gene set were computed by comparing our dataset to the permutated dataset for each single cell. We then plotted out the –log_10_(p-values) showing the enriched gene sets grouped by cell type or treatment using the ComplexHeatmap R package (V2.5.6)^97^. (**Figure 4I, 5C**)

### Construction of the lineage trajectory

We used Monocle3 (V0.2.0) and scVelo (V0.2.1) to identify the cell fate decision genes and pseudotime from expression matrix^57^. Monocle3 first projected cells onto a low-dimensional space using UMAP^58^, then grouped similar cells using the Louvain community detection algorithm, and constructed trajectories based on divergence or convergence of cells types (**Figure 3 F-G**). For RNA velocity, we first created a loom file of spliced and unspliced mRNAs from annotated sequencing bam file using Velocyto package (V0.17.16) in Python (V3.6)^98^. Then we performed calculations of cell dynamics using scVelo package (V0.2.1) in Python (V3.6)^48^ (**Figure 3 H-K, Figure 5 B**). The scVelo algorithm is based on RNA velocity, requiring minimal prior knowledge about the sample molecular signatures. It solves the full transcriptional dynamics and has been effectively tested in dissecting kinetics in neurogenesis.

### Identification of gene regulatory network

The Scenic package (V1.2.2) in R(V3.6.3) was used with default settings to identify key cell fate decision transcription networks.^99^ In brief, sets of genes that were co-expressed with transcription factors were identified using the GENIE3 module. We kept the top five transcription factors for each target gene. Then, the RcisTarget module sought out the enriched cis-regulatory motifs of candidate transcription factors and predicted the candidate target genes. Lastly, we used the AUCell algorithm in the SCENIC package to score the activity of each regulon in each cell. We grouped the scores by cell types and plotted their RegulonAUC values in the heatmaps to show the regulon activities of master regulators (**Figure 4A-B, 5A**, **Figure S5**). This method is robust against normalization methods and cell dropouts, as we got consistent results after testing multiple runs of the same datasets under different normalization pipelines.

## Sequencing Data Availability

The single-cell RNA sequencing data will be deposited in the Gene Expression Omnibus (GEO) following the acceptance of the manuscript.

## Supplemental Materials

Supplemental information includes 15 supplementary figures, 5 supplementary tables, 1 video, and supplemental text.

## Author Contributions

H. K., Y. X., B. X. and K. L. conceived the project and designed the experiments; H. K. Y. X., M. W., H. Z., S.C. and E. M. performed the experiments; Y. X., H. K, X. C., M. F., S. H., and J. I. analyzed the data; B. X. and K. L. supervised the project; H. K. and Y. X. wrote the paper.

## Acknowledgments

We thank Drs. Barbara Corneo, Peter Sims, Tingting Wu, and Dantong (Danielle) Huang for scientific discussion and suggestions. We thank Erin Bush for help on single cell sequencing. This research was supported by National Institutes of Health (UH3TR002151), National Research Foundation of Korea (GRL 2015K1A1A2032163) and Dankook University (Priority Institute Support Program in 2021, Global Research Program). Single cell sequencing was performed at Single Cell Analysis Core of Columbia Genome Center. Sequencing data was analyzed at Columbia high performance computing systems (Habanero and C2B2 computing clusters).

---

**Table S1.** Samples for single cell RNA sequencing.

| Sample ID | KH001 | KH002 | KH003 | KH004 | KH005 |
| --- | --- | --- | --- | --- | --- |
| Days in Culture | 53 | 53 | 77 | 77 | 77 + 2 |
| FTY treatment | untreated | acute (4Hr) | untreated | chronic (77 days) | chronic (77 days) withdrawal (2 days) |
| Sample name | D53_UT | D53_AC_FTY | D77_UT | D77_FTY | D79_WD |
| Cells sequenced by 10X chromium | 5131 | 5,499 | 5,252 | 4,781 | 4,847 |
| Mean reads per cell | 79,997 | 74,256 | 72,722 | 73,999 | 77,290 |
| Median genes per cell | 3,963 | 3,735 | 4,342 | 3,603 | 4,684 |
| Median UMI counts per Cell | 11,995 | 10,733 | 12,987 | 13,811 | 14,570 |
| Total genes | 25,960 | 25,772 | 25,316 | 24,780 | 25,090 |
| Total reads | 410,469,533 | 408,334,504 | 381,936,982 | 353,790,983 | 374,623,346 |

**Table S2.** Chemicals and materials.

| Name | Vendor | Catalog Number |
| --- | --- | --- |
| mTeSR Plus Basal Medium | Stemcell Technologies | 1000276 |
| ReLeSR | Stemcell Technologies | 05873 |
| StemPro™ Accutase™ Cell Dissociation Reagent | Gibco | A1110501 |
| Y-27632 dihydrochloride (ROCK Inhibitor) | Tocris Bioscience | 1254 |
| Matrigel GFR Membrane Matrix | Corning | 354230 |
| KnockOut™ DMEM/F-12 Medium | Gibco | 12660012 |
| KnockOut™ Serum Replacement (KOSR) | Gibco | 10828028 |
| Glutamax™ Supplement | Gibco | 35050061 |
| Minimum essential media-nonessential amino acids solution (MEM-NEAA) | Gibco | 11140050 |
| 2-mercaptoethanol | Gibco | 21985023 |
| Neurobasal™ Medium | Gibco | 21103049 |
| N-2 Supplement (100X) | Gibco | 17502048 |
| B-27™ Supplement (50X), minus Vitamin A | Gibco | 12587010 |
| LDN193189 | Stemgent | 040074 |
| SB431542 | Tocris | 1614 |
| Recombinant Mouse Sonic Hedgehog / Shh (C25II) N-Terminus | R&D Systems | 464-SH |
| Purmorphamine | Calbiochem | 540220 |
| CHIR99021 | Stemgent | 040004 |
| BDNF Human | ProSpec | CYT-207 |
| Ascorbic acid | Sigma | A-4034 |
| GDNF | ProSpec | CYT-305 |
| dibutyryl-cAMP | Calbiochem | 28745 |
| TGF-b3 | R&D Systems | 243-B3 |
| DAPT | Tocris | 2634 |

**Table S3.**
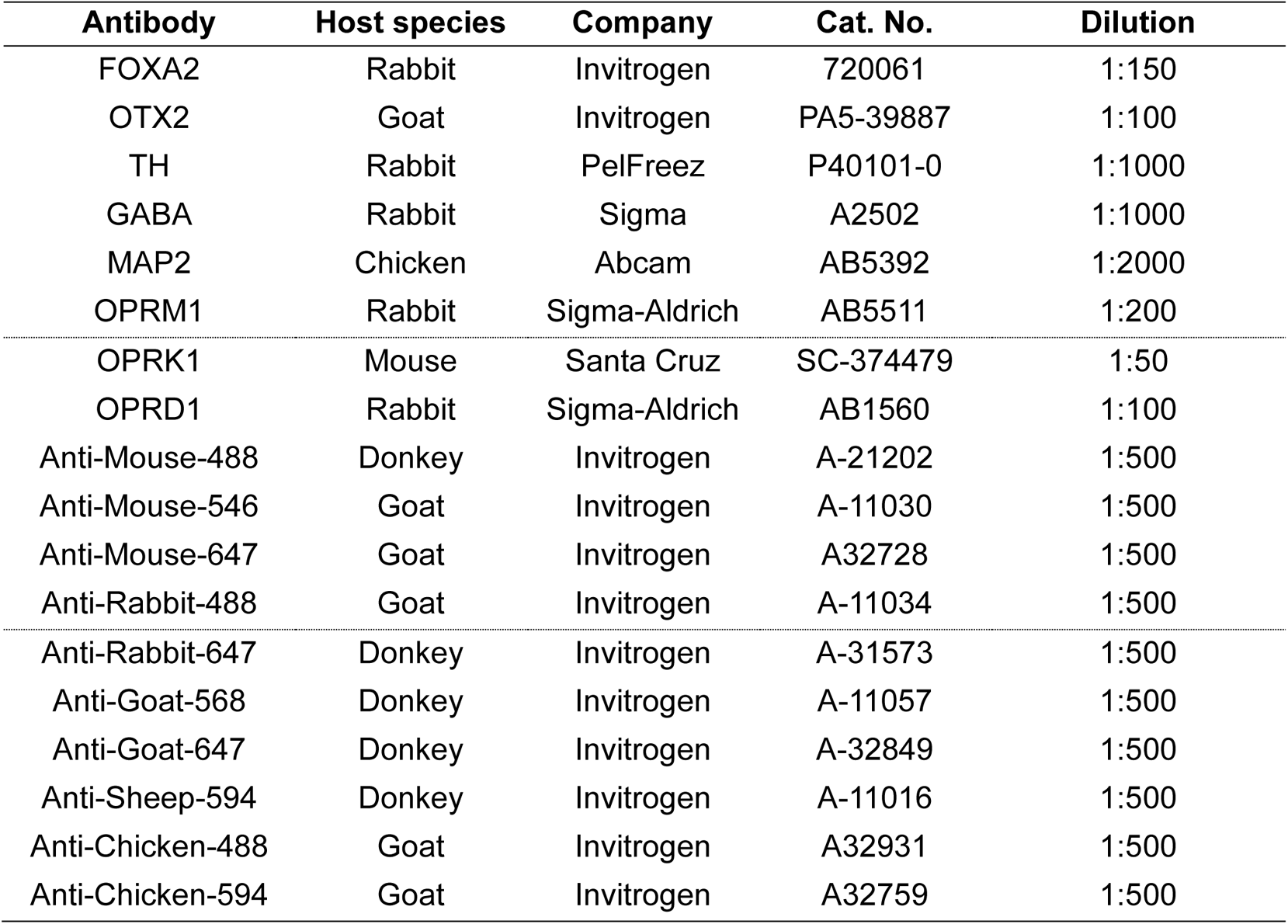
Antibodies.

**Table S4.**
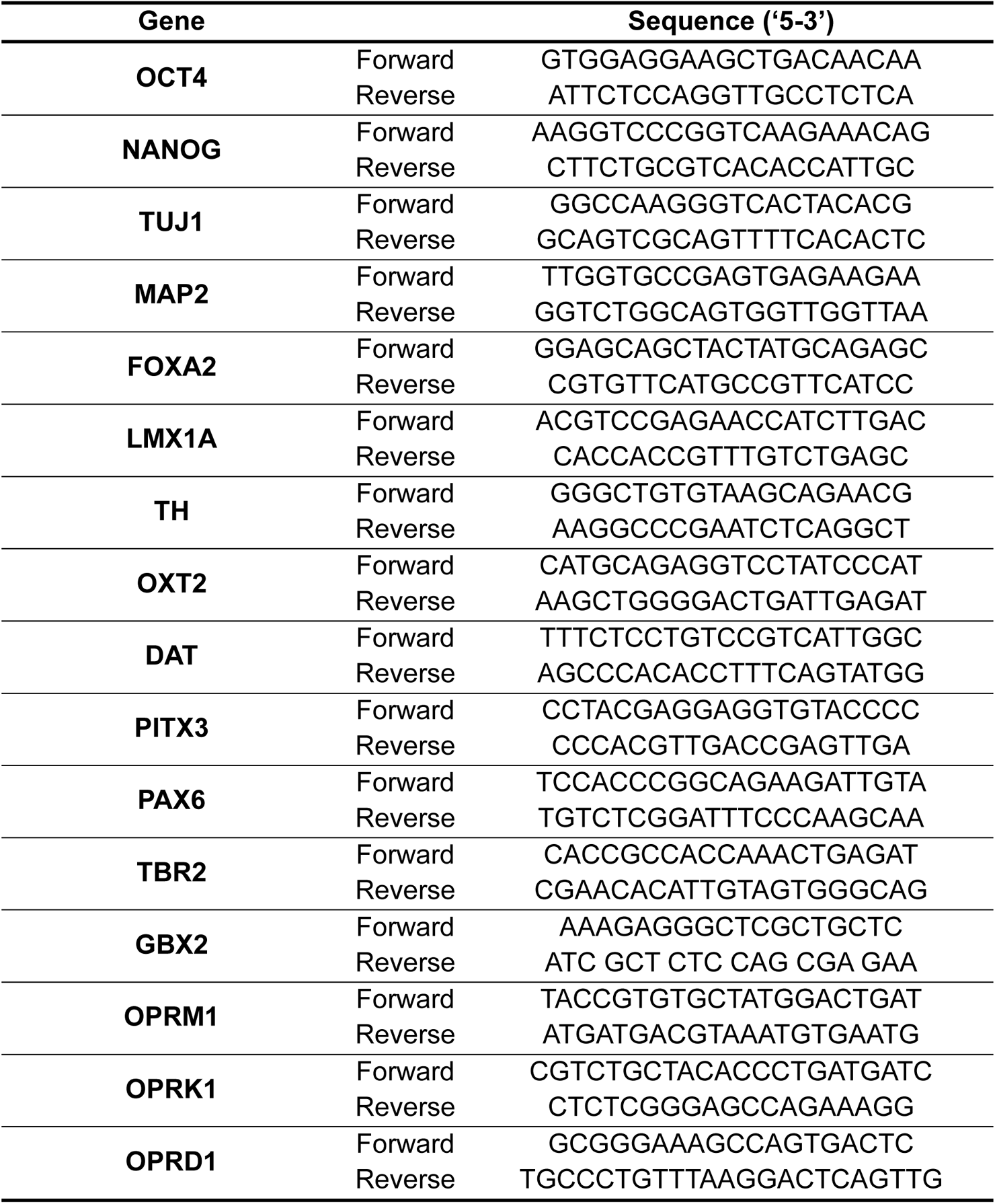
Primer sequences for real-time RT-PCR.

**Table S5.**
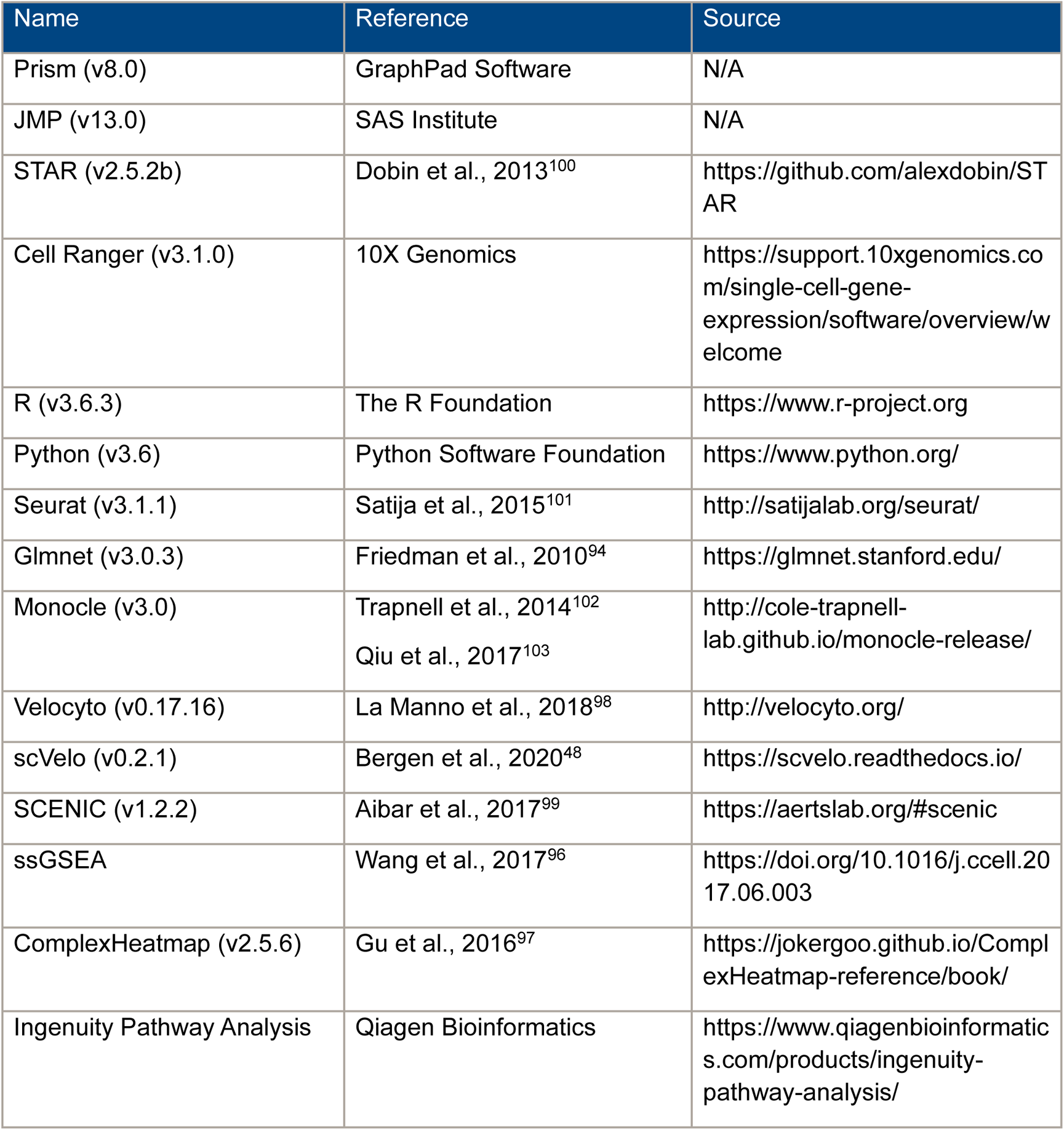
Software and algorithms.

## References

1. Sun Y, Bao Y, Kosten T, et al. Editorial: Challenges to Opioid Use Disorders During COVID-19. Am J Addict 2020; 29: 174–175. 2020/04/14. DOI: 10.1111/ajad.13031.

2. Hedegaard H, Bastian BA, Trinidad JP, et al. Regional differences in the drugs most frequently involved in drug overdose deaths: United States, 2017. 2019.

3. O’Donnell J, Gladden RM, Mattson CL, et al. Notes from the field: overdose deaths with carfentanil and other fentanyl analogs detected-10 states, July 2016–June 2017. Morbidity and Mortality Weekly Report 2018; 67: 767.

4. Spencer M, Warner M, Bastian BA, et al. Drug overdose deaths involving fentanyl, 2011–2016. 2019.

5. Wilson N, Kariisa M, Seth P, et al. Drug and opioid-involved overdose deaths-United States, 2017–2018. Morbidity and Mortality Weekly Report 2020; 69: 290–297.

6. Haley DF and Saitz R. The Opioid Epidemic During the COVID-19 Pandemic. JAMA 2020 2020/09/19. DOI: 10.1001/jama.2020.18543.

7. Niles JK, Gudin J, Radcliff J, et al. The Opioid Epidemic Within the COVID-19 Pandemic: Drug Testing in 2020. Popul Health Manag 2021; 24: S43–S51. 2020/10/09. DOI: 10.1089/pop.2020.0230.

8. Honein MA, Boyle C and Redfield RR. Public Health Surveillance of Prenatal Opioid Exposure in Mothers and Infants. Pediatrics 2019; 143 2019/01/19. DOI: 10.1542/peds.2018-3801.

9. Jantzie LL, Maxwell JR, Newville JC, et al. Prenatal opioid exposure: The next neonatal neuroinflammatory disease. Brain Behav Immun 2020; 84: 45–58. 2019/11/26. DOI: 10.1016/j.bbi.2019.11.007.

10. Ko JY, D’Angelo DV, Haight SC, et al. Vital Signs: Prescription Opioid Pain Reliever Use During Pregnancy - 34 U.S. Jurisdictions, 2019. MMWR Morb Mortal Wkly Rep 2020; 69: 897–903. 2020/07/17. DOI: 10.15585/mmwr.mm6928a1.

11. Patrick SW, Dudley J, Martin PR, et al. Prescription opioid epidemic and infant outcomes. Pediatrics 2015; 135: 842–850. DOI: 10.1542/peds.2014-3299.

12. Pryor JR, Maalouf FI, Krans EE, et al. The opioid epidemic and neonatal abstinence syndrome in the USA: a review of the continuum of care. Arch Dis Child Fetal Neonatal Ed 2017; 102: F183–F187. DOI: 10.1136/archdischild-2015-310045.

13. Camden A, Ray JG, To T, et al. Identification of Prenatal Opioid Exposure Within Health Administrative Databases. Pediatrics 2021; 147 2020/12/31. DOI: 10.1542/peds.2020-018507.

14. Doberczak TM, Thornton JC, Bernstein J, et al. Impact of maternal drug dependency on birth weight and head circumference of offspring. Am J Dis Child 1987; 141: 1163–1167. DOI: 10.1001/archpedi.1987.04460110033016.

15. Walhovd KB, Moe V, Slinning K, et al. Volumetric cerebral characteristics of children exposed to opiates and other substances in utero. Neuroimage 2007; 36: 1331–1344. DOI: 10.1016/j.neuroimage.2007.03.070.

16. Kotelchuck M, Cheng ER, Belanoff C, et al. The prevalence and impact of substance use disorder and treatment on maternal obstetric experiences and birth outcomes among singleton deliveries in Massachusetts. Maternal and child health journal 2017; 21: 893–902.

17. Lammers EM, Johnson PN, Ernst KD, et al. Association of fentanyl with neurodevelopmental outcomes in very-low-birth-weight infants. Annals of Pharmacotherapy 2014; 48: 335–342.

18. Attarian S, Tran LC, Moore A, et al. The neurodevelopmental impact of neonatal morphine administration. Brain Sci 2014; 4: 321–334. DOI: 10.3390/brainsci4020321.

19. Larson JJ, Graham DL, Singer LT, et al. Cognitive and Behavioral Impact on Children Exposed to Opioids During Pregnancy. Pediatrics 2019; 144. DOI: 10.1542/peds.2019-0514.

20. Conradt E, Crowell SE and Lester BM. Early life stress and environmental influences on the neurodevelopment of children with prenatal opioid exposure. Neurobiol Stress 2018; 9: 48–54. DOI: 10.1016/j.ynstr.2018.08.005.

21. Unger A, Metz V and Fischer G. Opioid dependent and pregnant: what are the best options for mothers and neonates? Obstet Gynecol Int 2012; 2012: 195954. 2012/04/13. DOI: 10.1155/2012/195954.

22. Winklbaur B, Kopf N, Ebner N, et al. Treating pregnant women dependent on opioids is not the same as treating pregnancy and opioid dependence: a knowledge synthesis for better treatment for women and neonates. Addiction 2008; 103: 1429–1440. 2008/09/12. DOI: 10.1111/j.1360-0443.2008.02283.x.

23. Byrnes EM and Vassoler FM. Modeling prenatal opioid exposure in animals: Current findings and future directions. Front Neuroendocrinol 2018; 51: 1–13. 2017/10/03. DOI: 10.1016/j.yfrne.2017.09.001.

24. Pankevich DE, Altevogt BM, Dunlop J, et al. Improving and accelerating drug development for nervous system disorders. Neuron 2014; 84: 546–553. 2014/12/03. DOI: 10.1016/j.neuron.2014.10.007.

25. Boggess T and Risher WC. Clinical and basic research investigations into the long-term effects of prenatal opioid exposure on brain development. J Neurosci Res 2020 2020/05/28. DOI: 10.1002/jnr.24642.

26. Radhakrishnan R, Grecco G, Stolze K, et al. Neuroimaging in infants with prenatal opioid exposure: Current evidence, recent developments and targets for future research. J Neuroradiol 2021; 48: 112–120. 2020/10/17. DOI: 10.1016/j.neurad.2020.09.009.

27. Saad MH, Rumschlag M, Guerra MH, et al. Differentially expressed gene networks, biomarkers, long noncoding RNAs, and shared responses with cocaine identified in the midbrains of human opioid abusers. Sci Rep 2019; 9: 1534. 2019/02/09. DOI: 10.1038/s41598-018-38209-8.

28. Bannon MJ, Johnson MM, Michelhaugh SK, et al. A molecular profile of cocaine abuse includes the differential expression of genes that regulate transcription, chromatin, and dopamine cell phenotype. Neuropsychopharmacology 2014; 39: 2191–2199. 2014/03/20. DOI: 10.1038/npp.2014.70.

29. Farris SP, Arasappan D, Hunicke-Smith S, et al. Transcriptome organization for chronic alcohol abuse in human brain. Mol Psychiatry 2015; 20: 1438–1447. 2014/12/03. DOI: 10.1038/mp.2014.159.

30. Zhou Z, Yuan Q, Mash DC, et al. Substance-specific and shared transcription and epigenetic changes in the human hippocampus chronically exposed to cocaine and alcohol. Proc Natl Acad Sci U S A 2011; 108: 6626–6631. 2011/04/06. DOI: 10.1073/pnas.1018514108.

31. Blondel D and Lutolf MP. Bioinspired Hydrogels for 3D Organoid Culture. Chimia (Aarau) 2019; 73: 81–85. 2019/03/01. DOI: 10.2533/chimia.2019.81.

32. Clevers H. Modeling Development and Disease with Organoids. Cell 2016; 165: 1586–1597. 2016/06/18. DOI: 10.1016/j.cell.2016.05.082.

33. Di Lullo E and Kriegstein AR. The use of brain organoids to investigate neural development and disease. Nat Rev Neurosci 2017; 18: 573–584. 2017/09/08. DOI: 10.1038/nrn.2017.107.

34. Qian X, Song H and Ming GL. Brain organoids: advances, applications and challenges. Development 2019; 146 2019/04/18. DOI: 10.1242/dev.166074.

35. Willner MJ, Xiao Y, Kim HS, et al. Modeling SARS-CoV-2 infection in individuals with opioid use disorder with brain organoids. J Tissue Eng 2021; 12: 2041731420985299. 2021/03/20. DOI: 10.1177/2041731420985299.

36. Jo J, Xiao Y, Sun Alfred X, et al. Midbrain-like Organoids from Human Pluripotent Stem Cells Contain Functional Dopaminergic and Neuromelanin-Producing Neurons. Cell Stem Cell 2016; 19: 248–257. DOI: 10.1016/j.stem.2016.07.005.

37. Kriks S, Shim J-W, Piao J, et al. Dopamine neurons derived from human ES cells efficiently engraft in animal models of Parkinson’s disease. Nature 2011; 480: 547–551. DOI: 10.1038/nature10648.

38. Bhaduri A, Andrews MG, Mancia Leon W, et al. Cell stress in cortical organoids impairs molecular subtype specification. Nature 2020; 578: 142–148. DOI: 10.1038/s41586-020-1962-0.

39. Pollen AA, Bhaduri A, Andrews MG, et al. Establishing Cerebral Organoids as Models of Human-Specific Brain Evolution. Cell 2019; 176: 743–756.e717. DOI: 10.1016/j.cell.2019.01.017.

40. Miura Y, Li MY, Birey F, et al. Generation of human striatal organoids and cortico-striatal assembloids from human pluripotent stem cells. Nat Biotechnol 2020; 38: 1421–1430. 2020/12/05. DOI: 10.1038/s41587-020-00763-w.

41. Pasca AM, Sloan SA, Clarke LE, et al. Functional cortical neurons and astrocytes from human pluripotent stem cells in 3D culture. Nat Methods 2015; 12: 671–678. 2015/05/26. DOI: 10.1038/nmeth.3415.

42. Yoon SJ, Elahi LS, Pasca AM, et al. Reliability of human cortical organoid generation. Nat Methods 2019; 16: 75–78. DOI: 10.1038/s41592-018-0255-0.

43. Sakaguchi H, Kadoshima T, Soen M, et al. Generation of functional hippocampal neurons from self-organizing human embryonic stem cell-derived dorsomedial telencephalic tissue. Nat Commun 2015; 6: 8896. DOI: 10.1038/ncomms9896.

44. Amin ND and Pasca SP. Building Models of Brain Disorders with Three-Dimensional Organoids. Neuron 2018; 100: 389–405. 2018/10/26. DOI: 10.1016/j.neuron.2018.10.007.

45. Quadrato G, Nguyen T, Macosko EZ, et al. Cell diversity and network dynamics in photosensitive human brain organoids. Nature 2017; 545: 48–53. 2017/04/27. DOI: 10.1038/nature22047.

46. Tanaka Y, Cakir B, Xiang Y, et al. Synthetic Analyses of Single-Cell Transcriptomes from Multiple Brain Organoids and Fetal Brain. Cell Rep 2020; 30: 1682–1689 e1683. 2020/02/13. DOI: 10.1016/j.celrep.2020.01.038.

47. Camp JG, Badsha F, Florio M, et al. Human cerebral organoids recapitulate gene expression programs of fetal neocortex development. Proc Natl Acad Sci U S A 2015; 112: 15672–15677. 2015/12/09. DOI: 10.1073/pnas.1520760112.

48. Bergen V, Lange M, Peidli S, et al. Generalizing RNA velocity to transient cell states through dynamical modeling. Nat Biotech 2020: 1–7. DOI: 10.1038/s41587-020-0591-3.

49. Aibar S, González-Blas CB, Moerman T, et al. SCENIC: single-cell regulatory network inference and clustering. Nat Meth 2017; 14: 1083–1086. DOI: 10.1038/nmeth.4463.

50. Baizabal JM and Covarrubias L. The embryonic midbrain directs neuronal specification of embryonic stem cells at early stages of differentiation. Dev Biol 2009; 325: 49–59. DOI: 10.1016/j.ydbio.2008.09.024.

51. Puelles E, Acampora D, Lacroix E, et al. Otx dose-dependent integrated control of antero-posterior and dorso-ventral patterning of midbrain. Nat Neurosci 2003; 6: 453–460. 2003/03/26. DOI: 10.1038/nn1037.

52. La Manno G, Gyllborg D, Codeluppi S, et al. Molecular Diversity of Midbrain Development in Mouse, Human, and Stem Cells. Cell 2016; 167: 566–580 e519. 2016/10/08. DOI: 10.1016/j.cell.2016.09.027.

53. Al-Hasani R and Bruchas MR. Molecular mechanisms of opioid receptor-dependent signaling and behavior. Anesthesiology 2011; 115: 1363–1381. 2011/10/25. DOI: 10.1097/ALN.0b013e318238bba6.

54. Karagiannis TT, Cleary JP, Jr., Gok B, et al. Single cell transcriptomics reveals opioid usage evokes widespread suppression of antiviral gene program. Nat Commun 2020; 11: 2611. 2020/05/28. DOI: 10.1038/s41467-020-16159-y.

55. Merhar SL, Kline JE, Braimah A, et al. Prenatal opioid exposure is associated with smaller brain volumes in multiple regions. Pediatr Res 2020 2020/11/13. DOI: 10.1038/s41390-020-01265-w.

56. Merhar SL, McAllister JM, Wedig-Stevie KE, et al. Retrospective review of neurodevelopmental outcomes in infants treated for neonatal abstinence syndrome. J Perinatol 2018; 38: 587–592. 2018/03/09. DOI: 10.1038/s41372-018-0088-9.

57. Cao J, Spielmann M, Qiu X, et al. The single-cell transcriptional landscape of mammalian organogenesis. Nature 2019; 566: 496–502. 2019/02/23. DOI: 10.1038/s41586-019-0969-x.

58. McInnes L, Healy J and Melville J. UMAP: Uniform Manifold Approximation and Projection for Dimension Reduction. *arXiv*:180203426 [*cs, stat*] 2020.

59. Eze UC, Bhaduri A, Nowakowski TJ, et al. Heterogeneity of Human Neuroepithelial Cells and Early Radial Glia. bioRxiv 2020: 2020.2003.2006.981423. DOI: 10.1101/2020.03.06.981423.

60. Mantamadiotis T, Papalexis N and Dworkin S. CREB signalling in neural stem/progenitor cells: recent developments and the implications for brain tumour biology. Bioessays 2012; 34: 293–300. 2012/02/15. DOI: 10.1002/bies.201100133.

61. Hecker N, Seemann SE, Silahtaroglu A, et al. Associating transcription factors and conserved RNA structures with gene regulation in the human brain. Sci Rep 2017; 7: 5776. 2017/07/20. DOI: 10.1038/s41598-017-06200-4.

62. Ding Q, Joshi PS, Xie ZH, et al. BARHL2 transcription factor regulates the ipsilateral/contralateral subtype divergence in postmitotic dI1 neurons of the developing spinal cord. Proc Natl Acad Sci U S A 2012; 109: 1566–1571. 2012/02/07. DOI: 10.1073/pnas.1112392109.

63. Ding B, Cave JW, Dobner PR, et al. Reciprocal autoregulation by NFI occupancy and ETV1 promotes the developmental expression of dendrite-synapse genes in cerebellar granule neurons. Mol Biol Cell 2016; 27: 1488–1499. 2016/03/05. DOI: 10.1091/mbc.E15-07-0476.

64. Abaci HE, Shen Y-I, Tan S, et al. Recapitulating physiological and pathological shear stress and oxygen to model vasculature in health and disease. Scientific Reports 2014; 4. DOI: 10.1038/srep04951.

65. Thakker-Varia S and Alder J. Neuropeptides in depression: role of VGF. Behav Brain Res 2009; 197: 262–278. 2008/11/06. DOI: 10.1016/j.bbr.2008.10.006.

66. Taylor AMW, Becker S, Schweinhardt P, et al. Mesolimbic dopamine signaling in acute and chronic pain: implications for motivation, analgesia, and addiction. Pain 2016; 157: 1194–1198. 2016/01/23. DOI: 10.1097/j.pain.0000000000000494.

67. Adinoff B. Neurobiologic processes in drug reward and addiction. Harv Rev Psychiatry 2004; 12: 305–320. 2005/03/15. DOI: 10.1080/10673220490910844.

68. Gordon A, Yoon SJ, Tran SS, et al. Long-term maturation of human cortical organoids matches key early postnatal transitions. Nat Neurosci 2021; 24: 331–342. DOI: 10.1038/s41593-021-00802-y.

69. Doucet-Beaupré H and Levesque M. The role of developmental transcription factors in adult midbrain dopaminergic neurons. OA Neurosciences 2013; 1: 3.

70. Birey F, Andersen J, Makinson CD, et al. Assembly of functionally integrated human forebrain spheroids. Nature 2017; 545: 54–59. DOI: 10.1038/nature22330.

71. Qian X, Nguyen HN, Song MM, et al. Brain-region-specific organoids using mini-bioreactors for modeling ZIKV exposure. Cell 2016; 165: 1238–1254.

72. Zhong S, Zhang S, Fan X, et al. A single-cell RNA-seq survey of the developmental landscape of the human prefrontal cortex. Nature 2018; 555: 524–528. DOI: 10.1038/nature25980.

73. Nellhaus EM, Murray S, Hansen Z, et al. Novel Withdrawal Symptoms of a Neonate Prenatally Exposed to a Fentanyl Analog. J Pediatr Health Care 2019; 33: 102–106. DOI: 10.1016/j.pedhc.2018.08.014.

74. Lepeta K, Lourenco MV, Schweitzer BC, et al. Synaptopathies: synaptic dysfunction in neurological disorders - A review from students to students. J Neurochem 2016; 138: 785–805. DOI: 10.1111/jnc.13713.

75. Quadrato G, Brown J and Arlotta P. The promises and challenges of human brain organoids as models of neuropsychiatric disease. Nat Med 2016; 22: 1220–1228. 2016/11/01. DOI: 10.1038/nm.4214.

76. Zhang DY, Song H and Ming GL. Modeling neurological disorders using brain organoids. Semin Cell Dev Biol 2020 2020/06/21. DOI: 10.1016/j.semcdb.2020.05.026.

77. Stoeckius M, Hafemeister C, Stephenson W, et al. Simultaneous epitope and transcriptome measurement in single cells. Nat Methods 2017; 14: 865–868. 2017/08/02. DOI: 10.1038/nmeth.4380.

78. Stickels RR, Murray E, Kumar P, et al. Highly sensitive spatial transcriptomics at near-cellular resolution with Slide-seqV2. Nat Biotech 2020: 1–7. DOI: 10.1038/s41587-020-0739-1.

79. Liu Y, Yang M, Deng Y, et al. High-Spatial-Resolution Multi-Omics Sequencing via Deterministic Barcoding in Tissue. Cell 2020; 183: 1665–1681 e1618. 2020/11/15. DOI: 10.1016/j.cell.2020.10.026.

80. Taber KH, Hurley RA and Yudofsky SC. Diagnosis and treatment of neuropsychiatric disorders. Annu Rev Med 2010; 61: 121–133. 2009/10/15. DOI: 10.1146/annurev.med.051408.105018.

81. Nock NL, Minnes S and Alberts JL. Neurobiology of substance use in adolescents and potential therapeutic effects of exercise for prevention and treatment of substance use disorders. Birth Defects Res 2017; 109: 1711–1729. 2017/12/19. DOI: 10.1002/bdr2.1182.

82. Ross S and Peselow E. The neurobiology of addictive disorders. Clin Neuropharmacol 2009; 32: 269–276. 2009/10/17. DOI: 10.1097/wnf.0b013e3181a9163c.

83. Patel A, Garcia Diaz A, Moore JC, et al. Establishment and characterization of two iPSC lines derived from healthy controls. Stem Cell Res 2020; 47: 101926. 2020/08/02. DOI: 10.1016/j.scr.2020.101926.

84. Chambers SM, Fasano CA, Papapetrou EP, et al. Highly efficient neural conversion of human ES and iPS cells by dual inhibition of SMAD signaling. Nat Biotechnol 2009; 27: 275–280. DOI: 10.1038/nbt.1529.

85. Joksimovic M, Yun BA, Kittappa R, et al. Wnt antagonism of Shh facilitates midbrain floor plate neurogenesis. Nat Neurosci 2009; 12: 125–131. DOI: 10.1038/nn.2243.

86. Kirkeby A, Grealish S, Wolf DA, et al. Generation of regionally specified neural progenitors and functional neurons from human embryonic stem cells under defined conditions. Cell Rep 2012; 1: 703–714. DOI: 10.1016/j.celrep.2012.04.009.

87. Chung SY, Kishinevsky S, Mazzulli JR, et al. Parkin and PINK1 Patient iPSC-Derived Midbrain Dopamine Neurons Exhibit Mitochondrial Dysfunction and alpha-Synuclein Accumulation. Stem Cell Reports 2016; 7: 664–677. 2016/09/20. DOI: 10.1016/j.stemcr.2016.08.012.

88. Lieberman OJ, Choi SJ, Kanter E, et al. alpha-Synuclein-Dependent Calcium Entry Underlies Differential Sensitivity of Cultured SN and VTA Dopaminergic Neurons to a Parkinsonian Neurotoxin. eNeuro 2017; 4 2017/11/28. DOI: 10.1523/ENEURO.0167-17.2017.

89. Woodard CM, Campos BA, Kuo SH, et al. iPSC-derived dopamine neurons reveal differences between monozygotic twins discordant for Parkinson’s disease. Cell Rep 2014; 9: 1173–1182. 2014/12/03. DOI: 10.1016/j.celrep.2014.10.023.

90. Mosharov EV, Staal RG, Bove J, et al. Alpha-synuclein overexpression increases cytosolic catecholamine concentration. J Neurosci 2006; 26: 9304–9311. 2006/09/08. DOI: 10.1523/JNEUROSCI.0519-06.2006.

91. Pothos E, Desmond M and Sulzer D. L-3,4-dihydroxyphenylalanine increases the quantal size of exocytotic dopamine release in vitro. J Neurochem 1996; 66: 629–636. 1996/02/01. DOI: 10.1046/j.1471-4159.1996.66020629.x.

92. Butler A, Hoffman P, Smibert P, et al. Integrating single-cell transcriptomic data across different conditions, technologies, and species. Nat Biotechnol 2018; 36: 411–420. 2018/04/03. DOI: 10.1038/nbt.4096.

93. Stuart T, Butler A, Hoffman P, et al. Comprehensive Integration of Single-Cell Data. Cell 2019; 177: 1888–1902 e1821. 2019/06/11. DOI: 10.1016/j.cell.2019.05.031.

94. Friedman J, Hastie T and Tibshirani R. Regularization Paths for Generalized Linear Models via Coordinate Descent. J Stat Softw 2010; 33: 1–22. 2010/09/03.

95. Simon N, Friedman J, Hastie T, et al. Regularization Paths for Cox’s Proportional Hazards Model via Coordinate Descent. J Stat Softw 2011; 39: 1–13. 2011/03/01. DOI: 10.18637/jss.v039.i05.

96. Wang Q, Hu B, Hu X, et al. Tumor Evolution of Glioma-Intrinsic Gene Expression Subtypes Associates with Immunological Changes in the Microenvironment. Cancer Cell 2017; 32: 42–56 e46. 2017/07/12. DOI: 10.1016/j.ccell.2017.06.003.

97. Gu Z, Eils R and Schlesner M. Complex heatmaps reveal patterns and correlations in multidimensional genomic data. Bioinformatics 2016; 32: 2847–2849. 2016/05/22. DOI: 10.1093/bioinformatics/btw313.

98. La Manno G, Soldatov R, Zeisel A, et al. RNA velocity of single cells. Nature 2018; 560: 494–498. 2018/08/10. DOI: 10.1038/s41586-018-0414-6.

99. Aibar S, Gonzalez-Blas CB, Moerman T, et al. SCENIC: single-cell regulatory network inference and clustering. Nat Methods 2017; 14: 1083–1086. 2017/10/11. DOI: 10.1038/nmeth.4463.

100. Dobin A, Davis CA, Schlesinger F, et al. STAR: ultrafast universal RNA-seq aligner. Bioinformatics 2013; 29: 15–21. 2012/10/30. DOI: 10.1093/bioinformatics/bts635.

101. Satija R, Farrell JA, Gennert D, et al. Spatial reconstruction of single-cell gene expression data. Nat Biotechnol 2015; 33: 495–502. 2015/04/14. DOI: 10.1038/nbt.3192.

102. Trapnell C, Cacchiarelli D, Grimsby J, et al. The dynamics and regulators of cell fate decisions are revealed by pseudotemporal ordering of single cells. Nat Biotechnol 2014; 32: 381–386. 2014/03/25. DOI: 10.1038/nbt.2859.

103. Qiu X, Mao Q, Tang Y, et al. Reversed graph embedding resolves complex single-cell trajectories. Nat Methods 2017; 14: 979–982. 2017/08/22. DOI: 10.1038/nmeth.4402.

