## Supplemental Information for "Chronic opioid treatment arrests neurodevelopment and alters synaptic activity in human midbrain organoids"

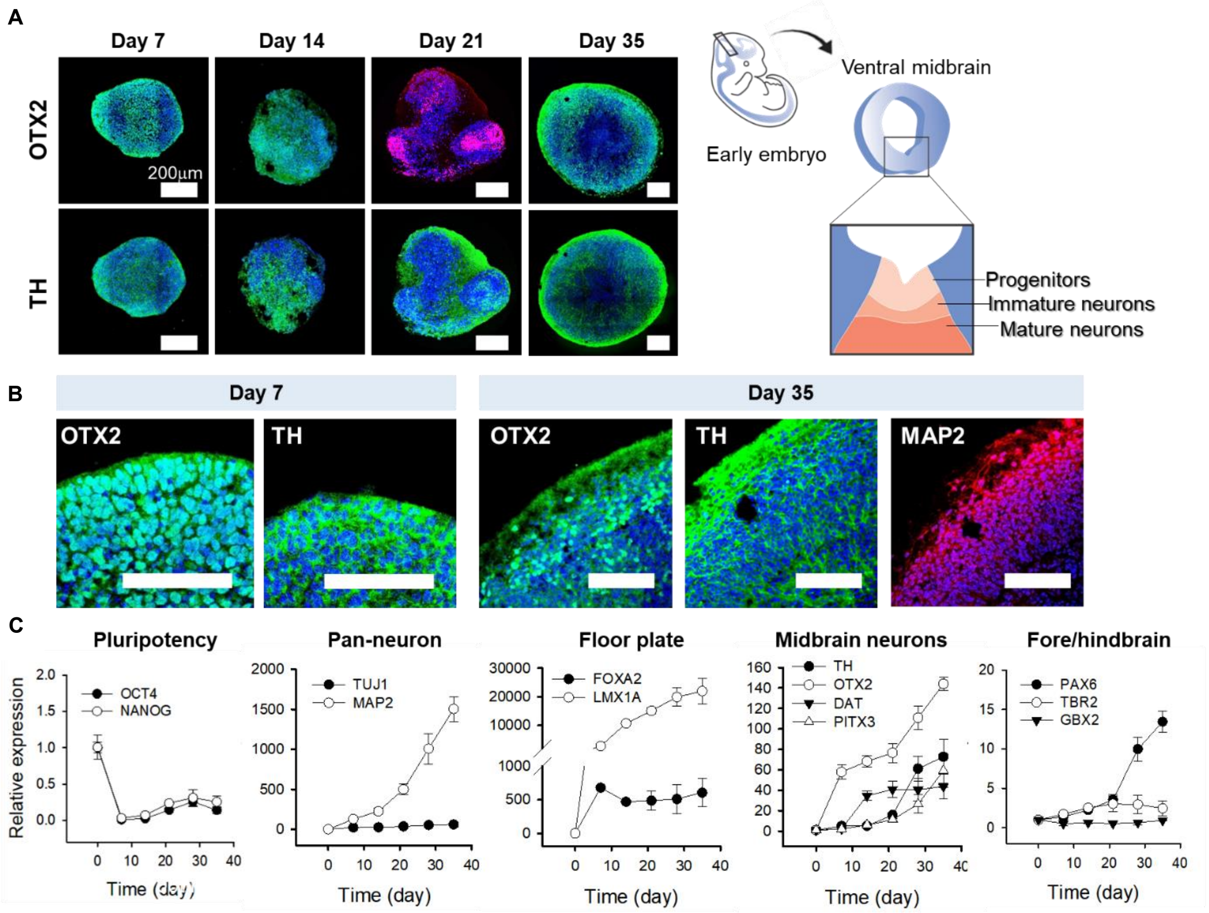


**Figure S1. Midbrain-regional development of organoids.**

(A-B) Immunocytochemical staining of the organoids for OTX2, TH, and MAP2. The unique zonal cytoarchitecture was clearly shown since day 21 of culture.

(C) Gene expression profiles of the organoids for the 35 day organoids by qRT-PCR (n=3-5 organoids).

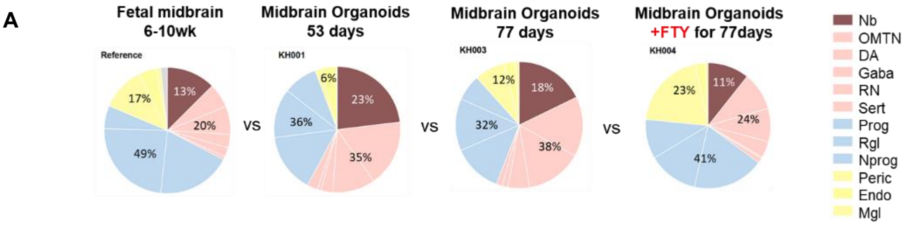


| Sample | Sample ID | Progenitors | | | | Neurons | | | | | Non-neurons | | | |
|---|---|---|---|---|---|---|---|---|---|---|---|---|---|---|
| | | Nb | Prog | Rgl | NProg | OMTN | DA | Gaba | RN | Sert | Peric | Endo | Mgl | OPC |
| 53d UT | KH001 | 23.09% | 15.13% | 12.65% | 8.75% | 17.14% | 10.83% | 3.06% | 0.95% | 2.70% | 1.42% | 4.01% | 0.26% | 0.00% |
| 77d UT | KH003 | 17.76% | 12.54% | 13.35% | 6.64% | 15.34% | 14.26% | 5.49% | 1.48% | 1.52% | 7.75% | 2.83% | 1.04% | 0.00% |
| 77d FTY | KH004 | 10.71% | 18.49% | 12.92% | 10.15% | 9.85% | 8.58% | 4.21% | 0.40% | 1.27% | 20.16% | 2.54% | 0.73% | 0.00% |
| Ref | Ref | 12.51% | 19.06% | 23.78% | 5.90% | 6.25% | 7.20% | 3.48% | 2.36% | 0.83% | 11.27% | 4.54% | 1.12% | 1.71% |

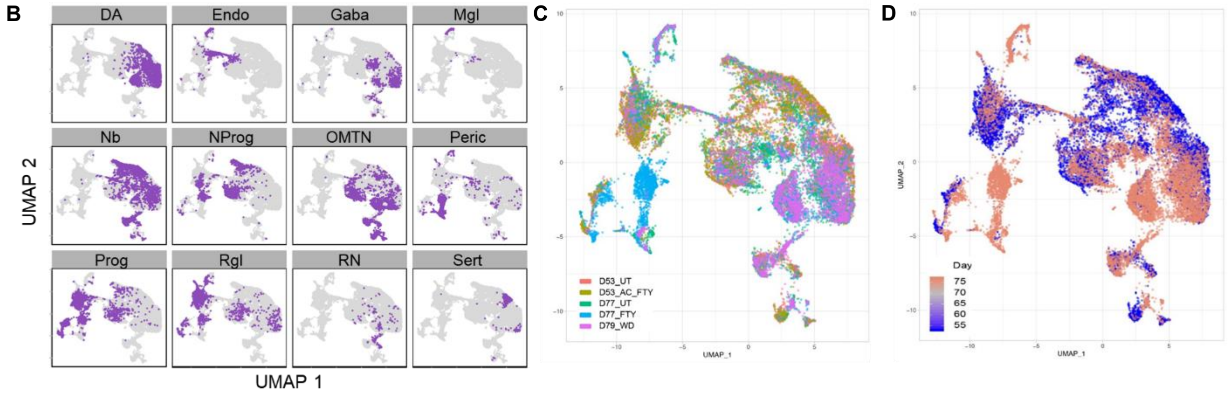


**Figure S2. Cell type identification and composition of the iPSC-derived midbrain-like organoids by scRNA seq.** The analysis was based on comparison to dataset of human fetal ventral midbrain of in vivo embryonic development at 6-10wk.

(A) Cell type compositions of midbrain organoids at day 53 or 77 of in vitro differentiation. (supplementary to Figure 1G).
(B) Cell types shown as monochrome plots in UMAP (supplementary to Figure 1F).
(C) Distribution in UMAP by sample (supplementary to Figure 1F).
(D) Distribution in UMAP by organoid age (supplementary to Figure 1F).

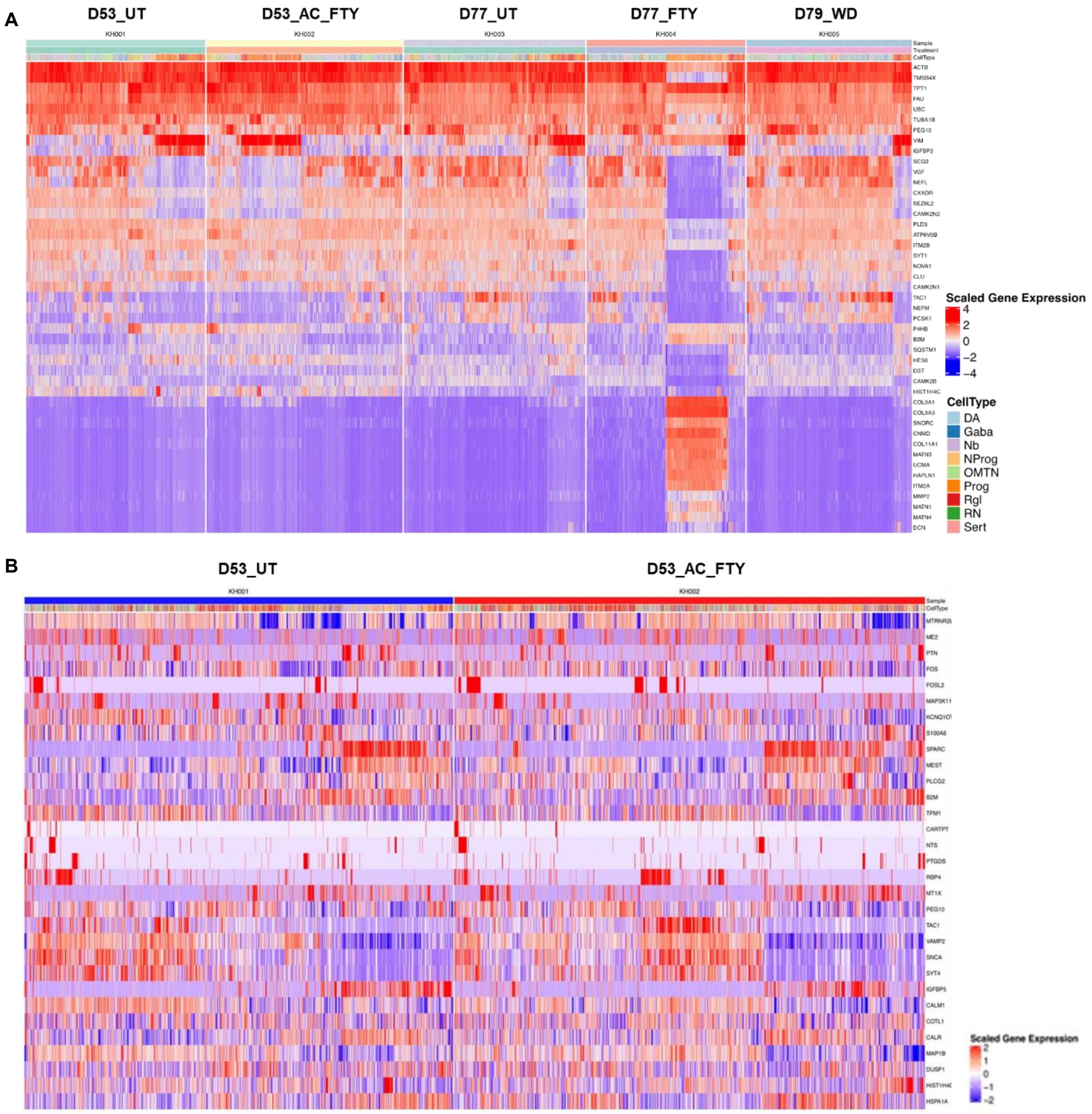


**Figure S3. Differentially expressed genes.**

(A) Top differentially expressed genes among all samples.

(B) Top differentially expressed genes detected in acute fentanyl treatment (comparing D53_UT vs D53_AC_FTY).

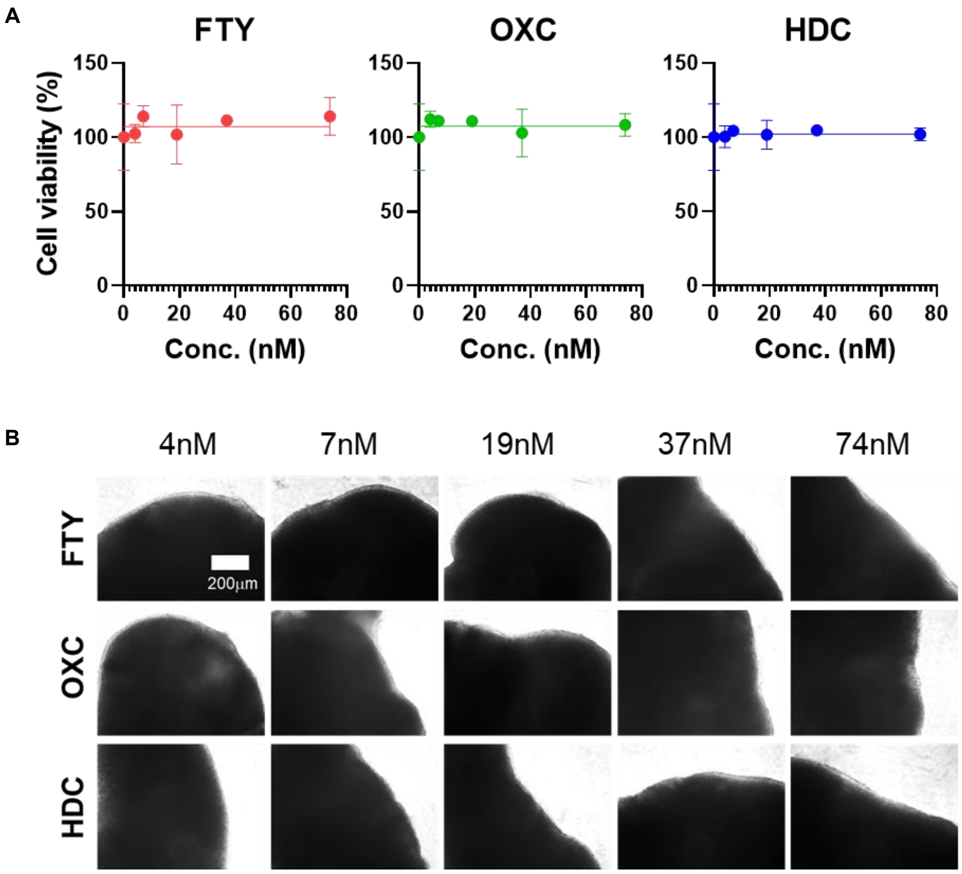


**Figure S4. Cell viability tests of the organoids with various opioid treatment (n=3 organoids, day 90 organoids).** (A) Cell viability depending on the concentration of opioids. (B) Bright-field images showing no noticeable dissociation of organoids.

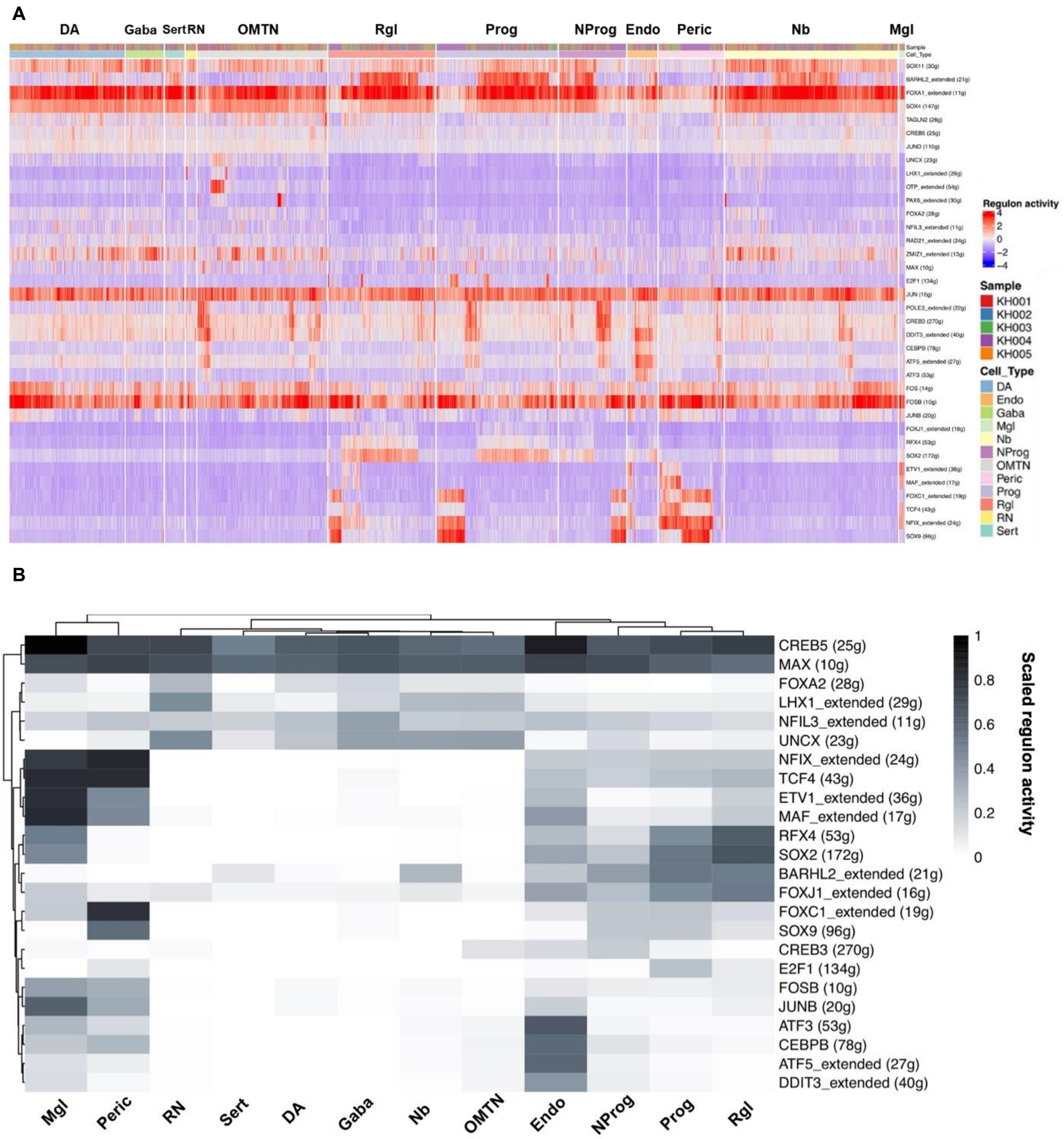


**Figure S5. Activities of gene regulatory network in all samples.**

(A) Master regulators identified in each single cell. Regulon activity is colored by scaled AUCell scores.
(B) Binarized regulon activity matrix by applying a cutoff minimum of 0.2 in AUCell score.

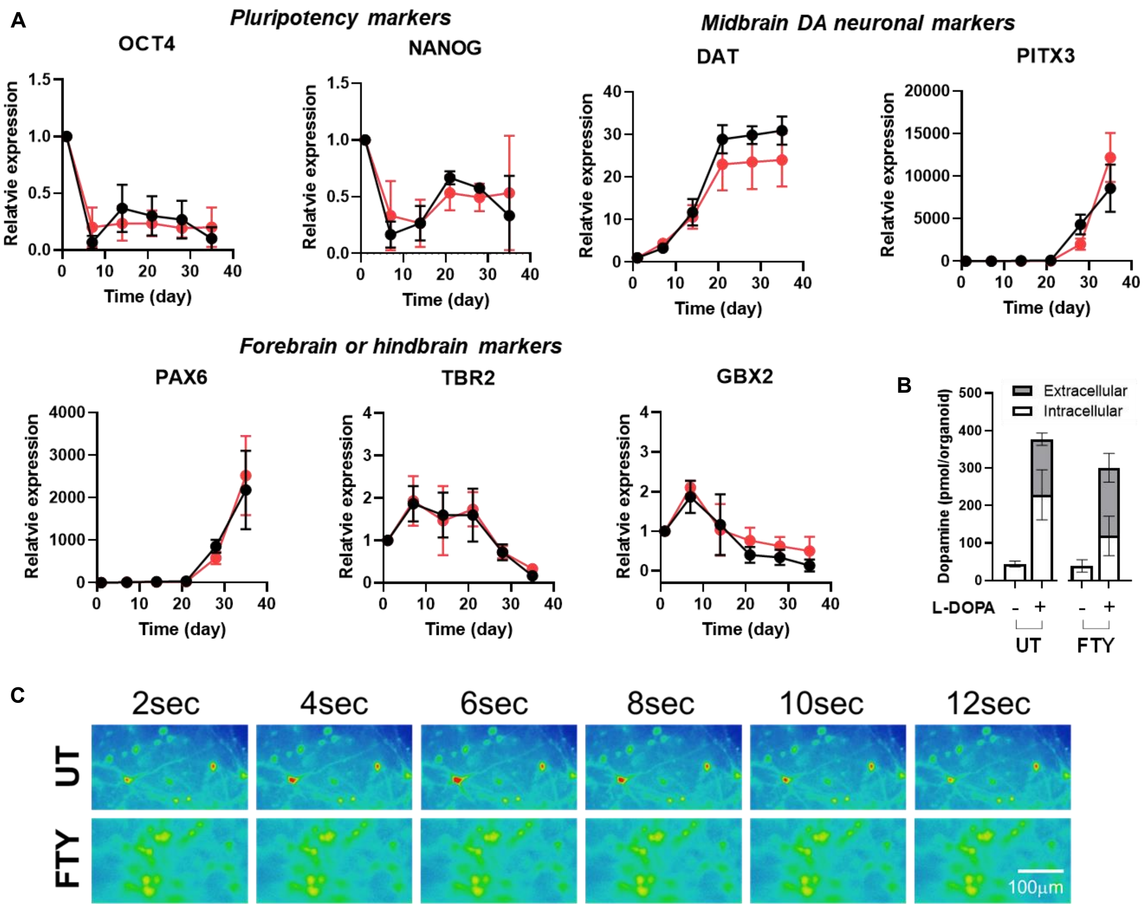


**Figure S6. Chronic opioid exposure on midbrain organoids and its development.**

(A) Fold changes of target genes of pluripotency gene (OCT4 and NANOG), midbrain dopaminergic neuronal genes (DAT and PITX3), forebrain or hindbrain genes (PAX6, TBR2, and GBX2) (n=5-7 organoids, three individual experiments).

(B) Dopamine synthesis and release of organoids with or without chronic fentanyl treatment (n=3 organoids, day 90 organoid).

(C) Calcium imaging of Day 180 organoids. Organoids with chronic fentanyl treatment showed no calcium influx (**Movie S2**).

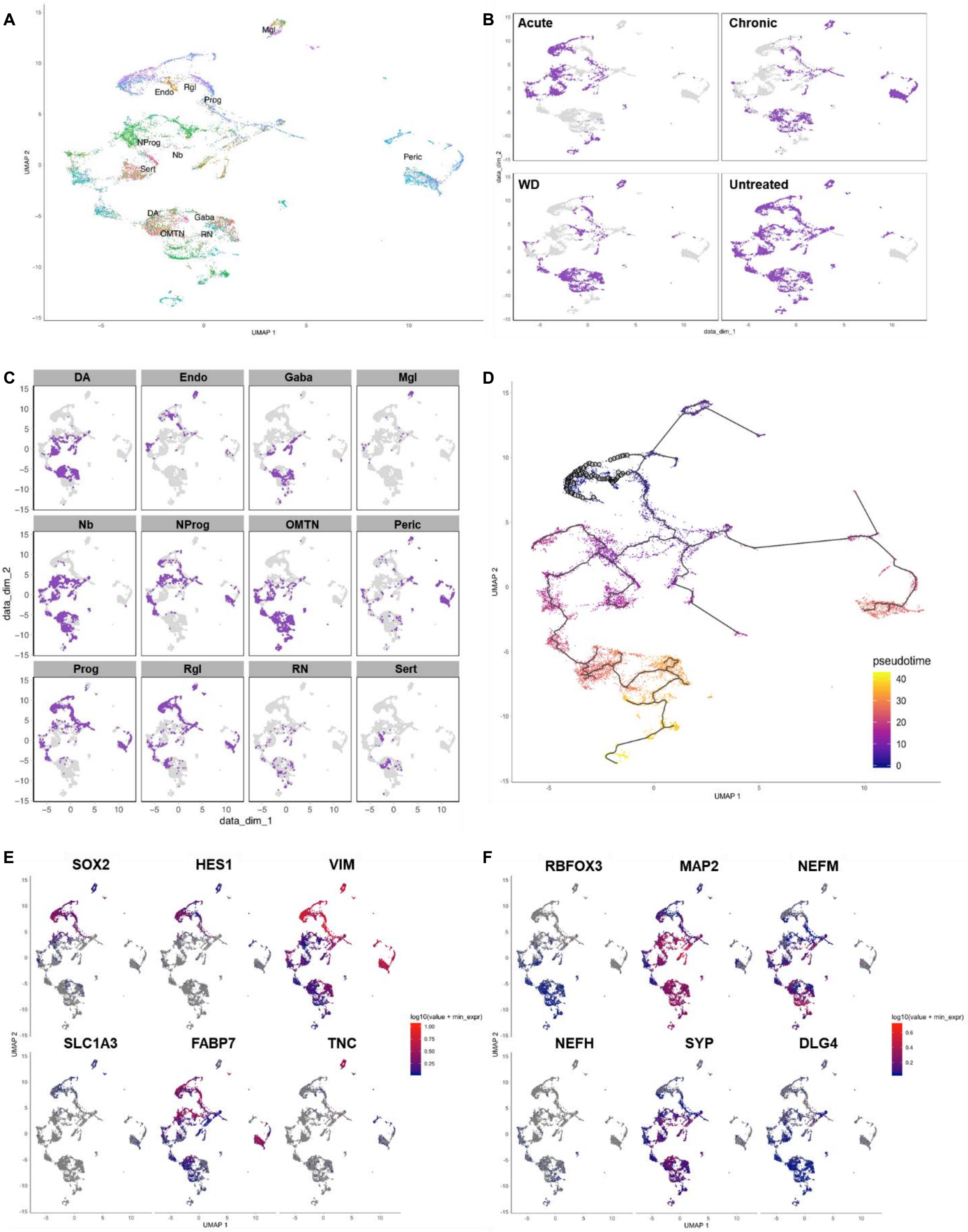


**Figure S7. Cell lineage trajectory analysis (Monocle) (Supplementary to Figure 3).**

(A) The Monocle plot showing the distribution of all samples by cell type.
(B) Monochrome plots showing the distribution of each sample.
(C) Monochrome plots showing each cell type.
(D) Lineage Trajectory in pseudotime.
(E) Feature plots of stem cell markers (SOX2, HES1, VIM, SLC1A3, FABP7, TNC).
(F) Feature plots of mature neuron markers (RBFOX3, MAP2, NEFM, NEFH, SYP, DLG4).

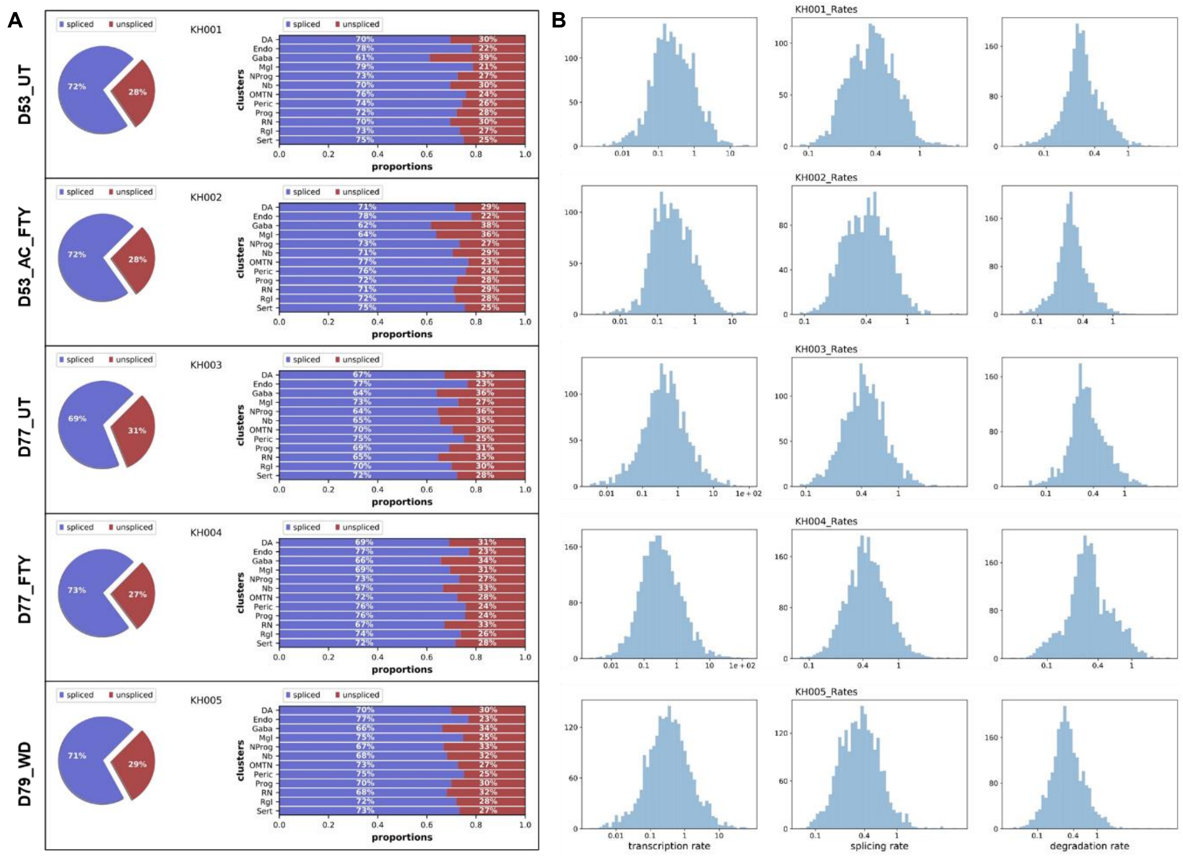

**Figure S8. Detailed plots of cell lineage trajectory analysis by RNA velocity. (Supplementary to Figure 3 H-K).**

(A) Proportions of spliced and unsliced mRNAs for all samples.

(B) Dynamic rates of RNA velocity in all five samples.

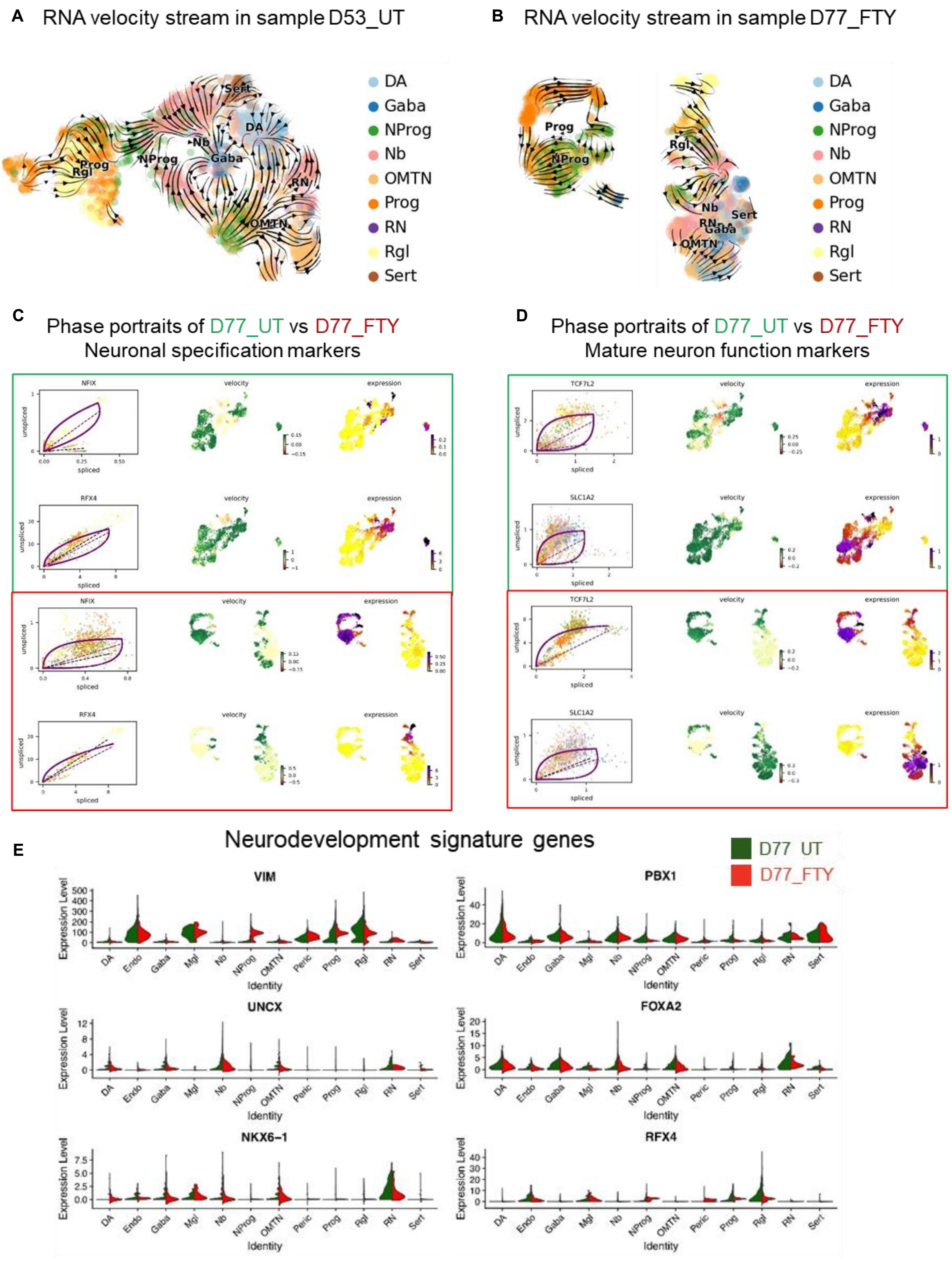


**Figure S9. Chronic Fentanyl treatment impairs neurodevelopment, shown by RNA velocity.**

A-B) RNA velocity computed based on spliced / unspliced mRNAs in D53_UT and D77_FTY sample.

C-D) Phase portraits of key genes in neuronal specification or mature neuron function.

E) Violin plots of signature genes in neurodevelopment.

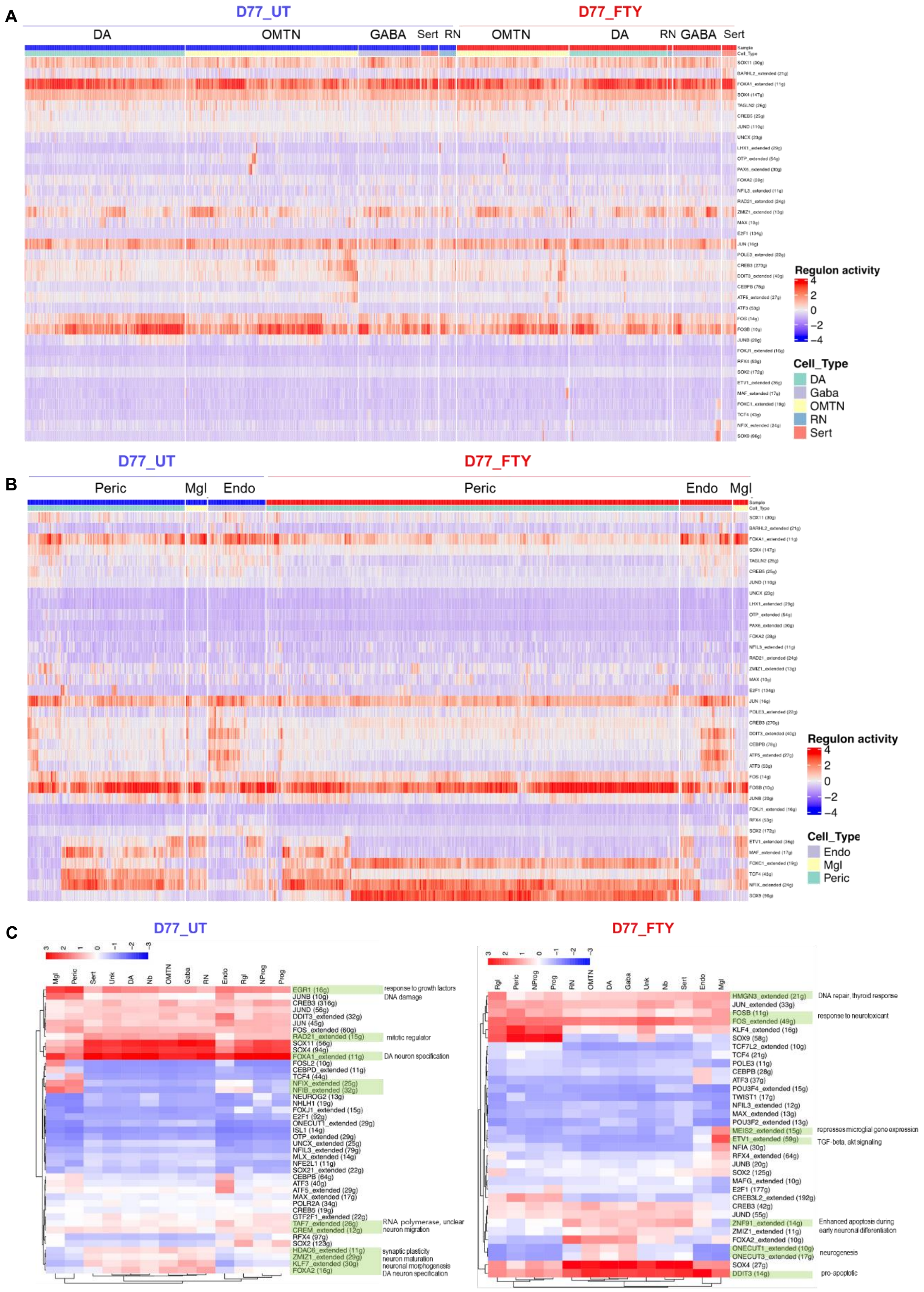


**Figure S10. Cell-type specific response to chronic fentanyl treatment, supplementary to Figure 4 (KH004 vs. KH003).** Critical Regulators of cell identity constructed by SCENIC R package.

(A) Master regulators identified in each single neuron. Regulon activity is colored by scaled AUCell scores.
(B) Master regulators identified in each mesoderm-derived cells.
(C) Master regulators identified in sample.

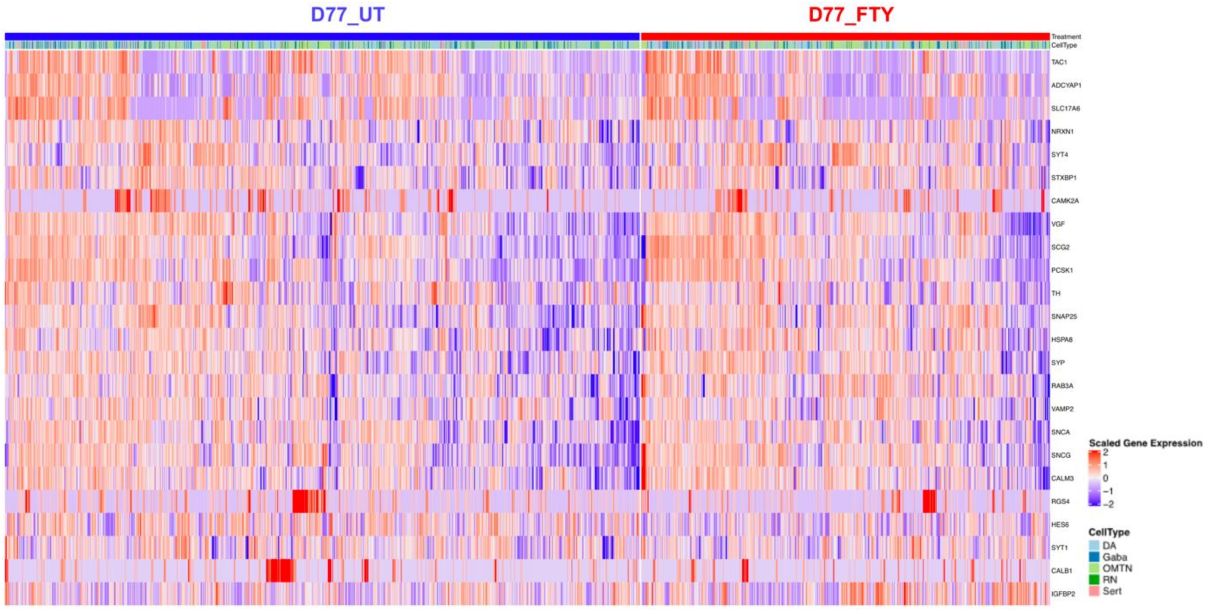


**Figure S11. Differentially expressed genes in neurons in response to chronic fentanyl treatment.**

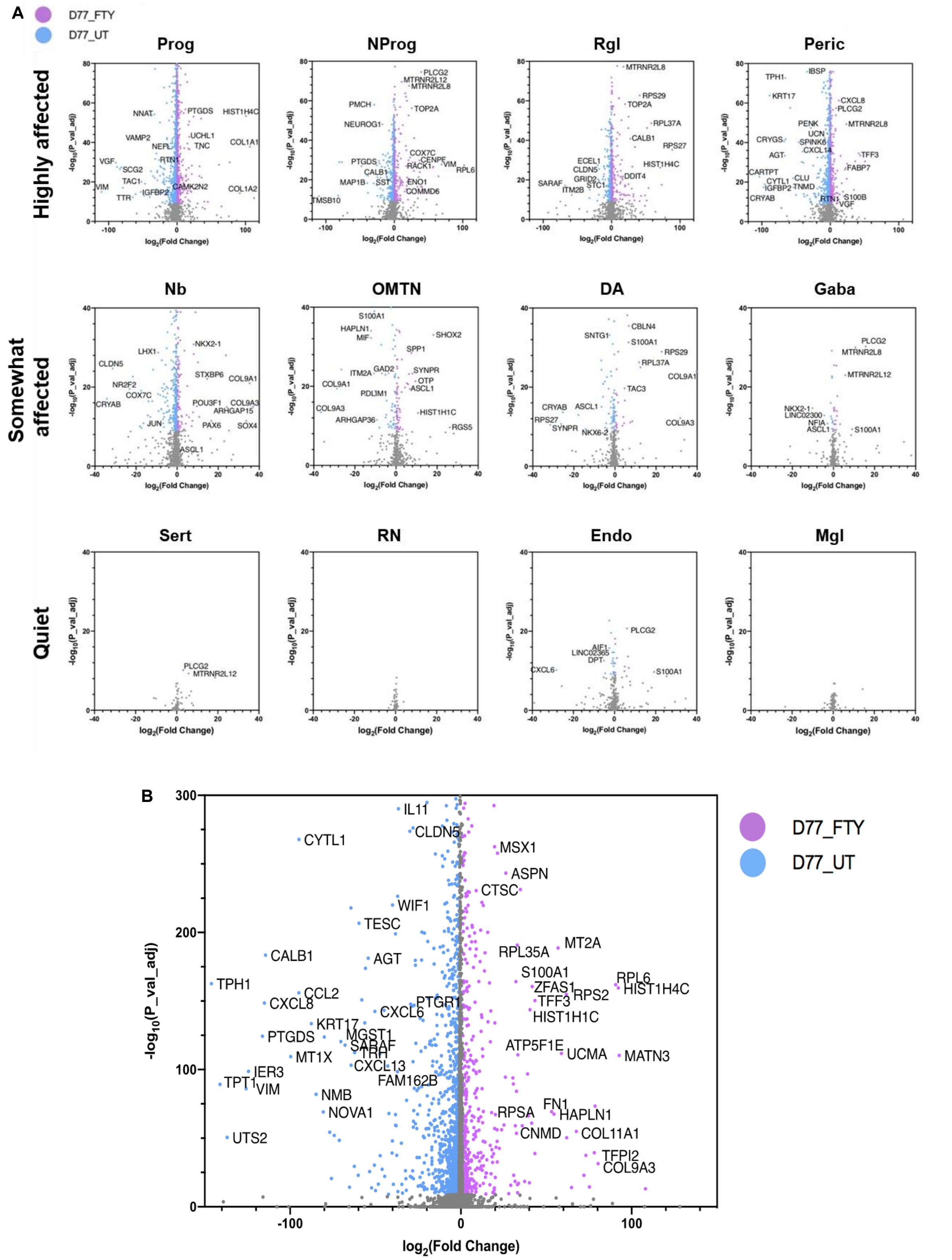


**Figure S12. Differential expression analysis showed cell-type specific response to chronic fentanyl treatment (Supplementary to Figure 4).**

(A) By cell type.

(B) Volcano plots of all cells in each sample.

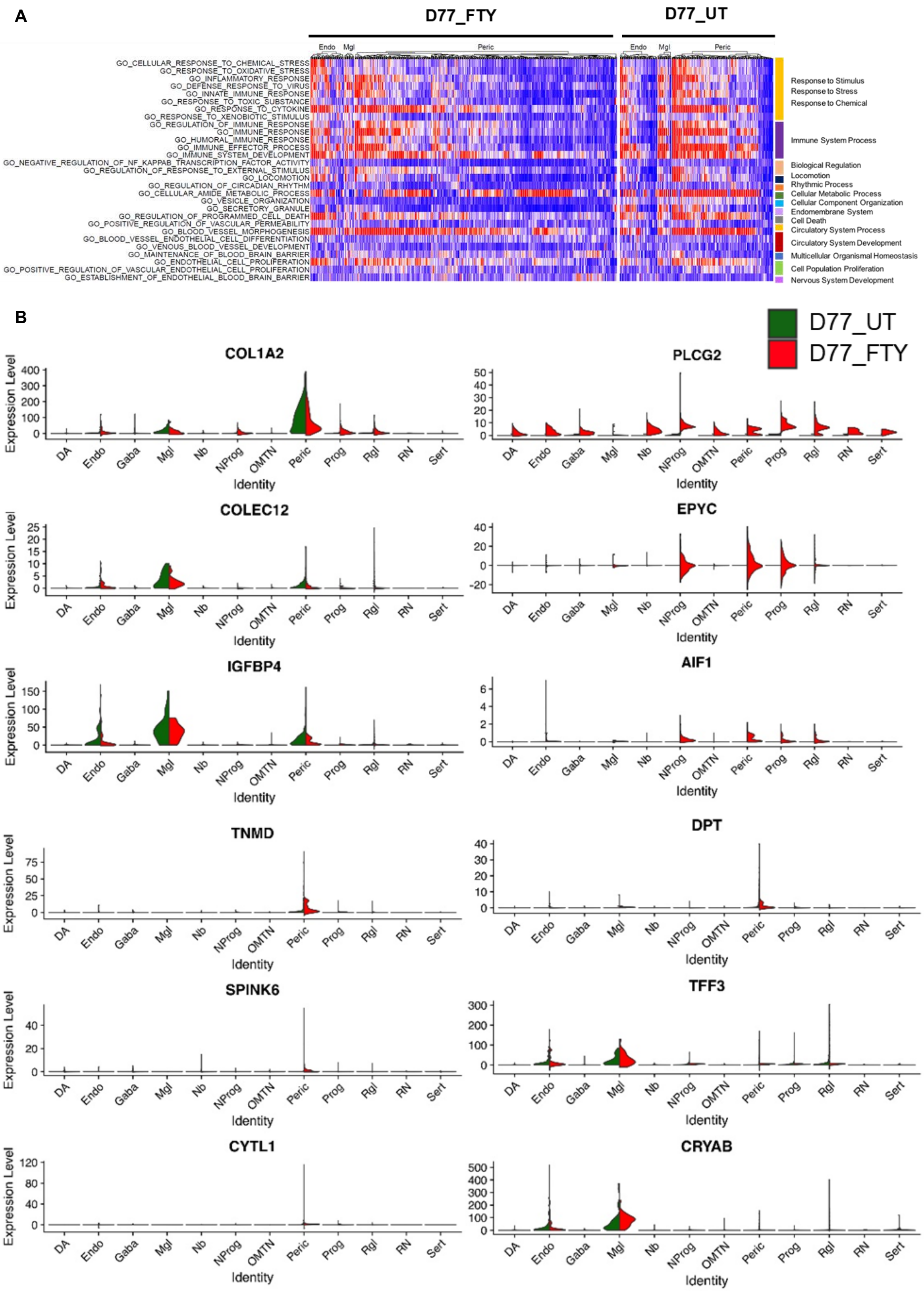


**Figure S13. Cell-type specific response to chronic fentanyl treatment (supplementary to Figure 4).**

(A) Mesoderm-derived cells; stimulus response and metabolic process

(B) Comparison of marker gene profiles in mesoderm-derived cells

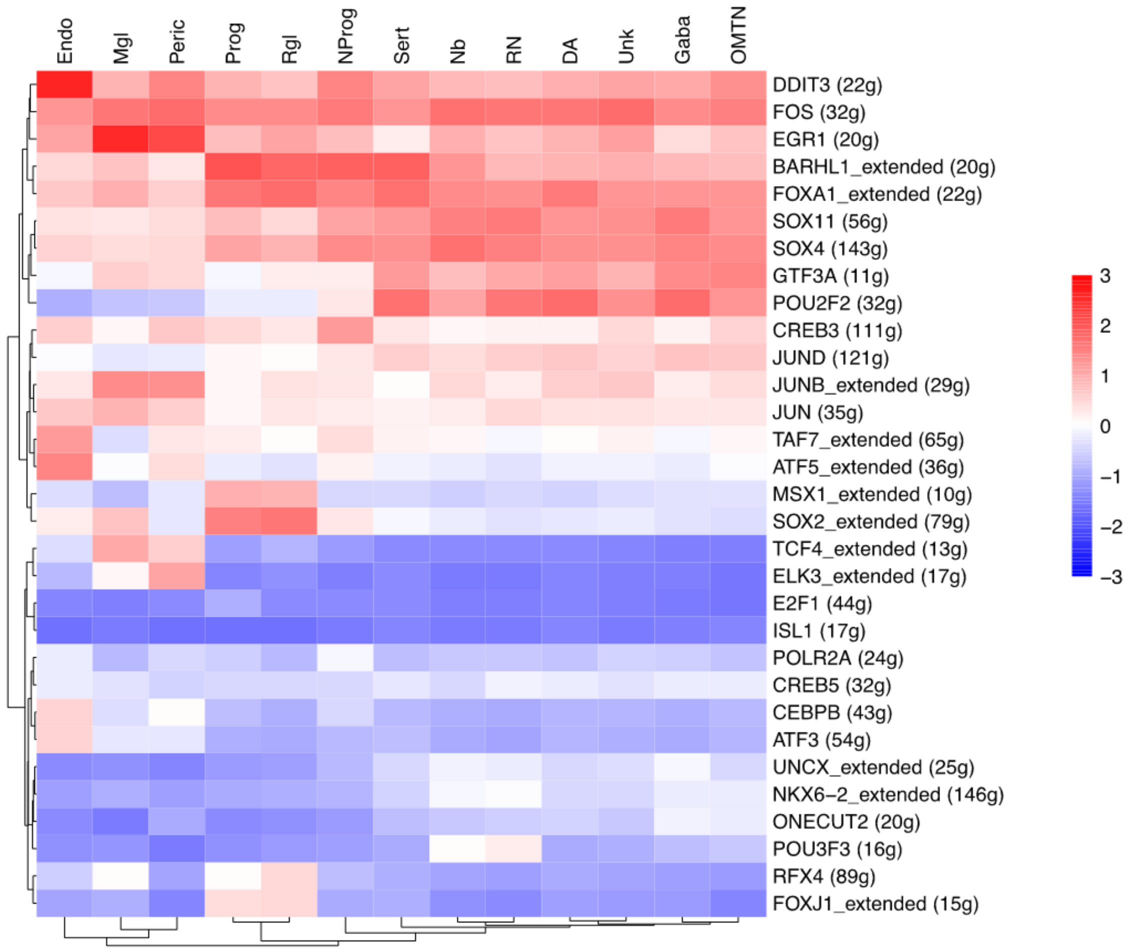


**Figure S14. Critical Regulators of cell identity constructed by SCENIC R Package (D79_WD).**

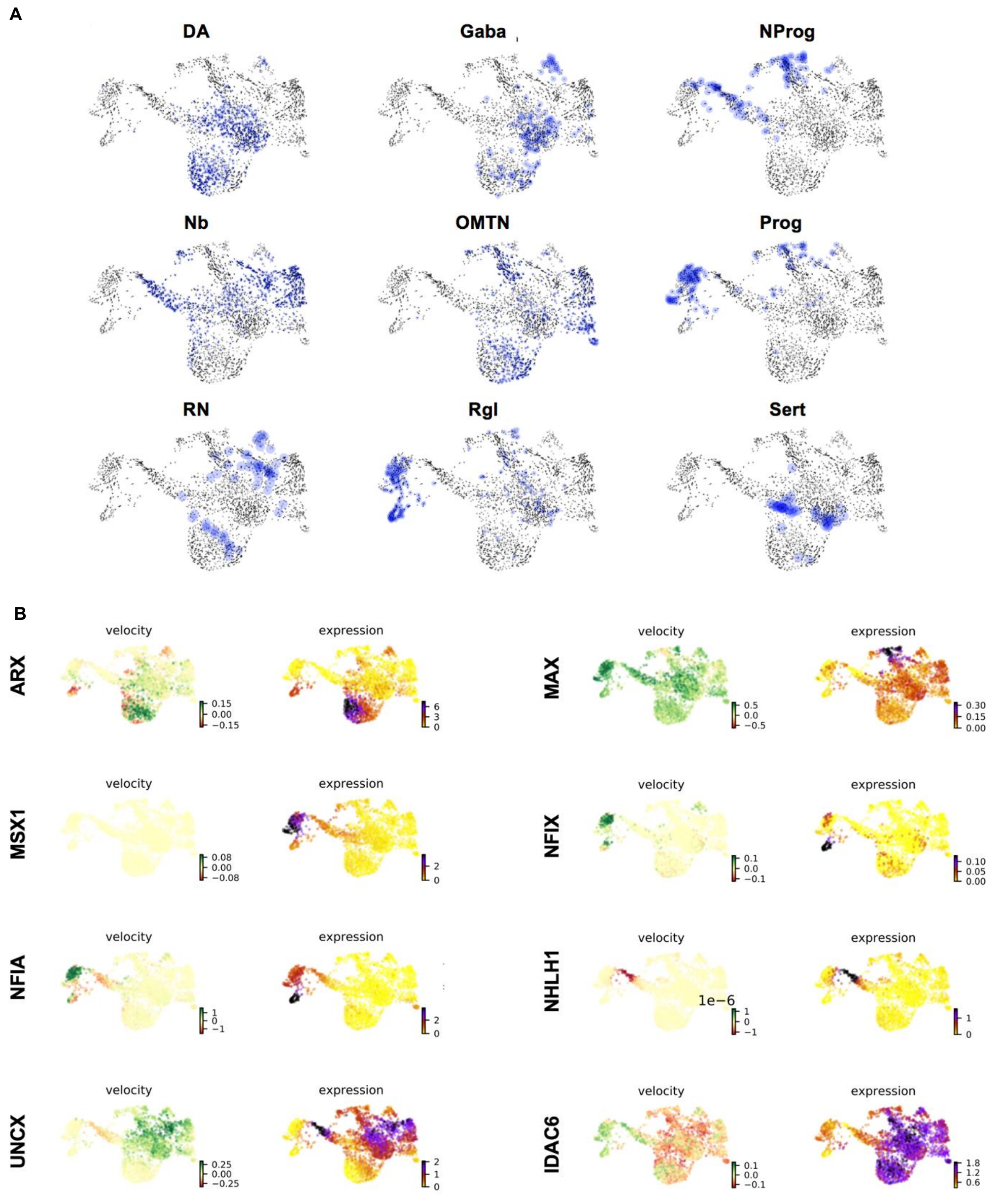


**Figure S15. RNA velocity (Supplementary to Figure 5).**

(A) Monochrome plots of cell types in D79_WD.
(B) Phase portraits of key genes in D79_WD.
